# Additive Dose Response Models: Defining Synergy

**DOI:** 10.1101/480608

**Authors:** Simone Lederer, Tjeerd M.H. Dijkstra, Tom Heskes

## Abstract

In synergy studies, one focuses on compound combinations that promise a synergistic or antagonistic effect. With the help of high-throughput techniques, a huge amount of compound combinations can be screened and filtered for suitable candidates for a more detailed analysis. Those promising candidates are chosen based on the deviance between a measured response and an expected non-interactive response. A non-interactive response is based on a principle of no interaction, such as Loewe Additivity [Loewe, 1928] or Bliss Independence [Bliss, 1939]. In Lederer et al. [2018a], an explicit formulation of the hitherto implicitly defined Loewe Additivity has been introduced, the so-called Explicit Mean Equation. In the current study we show that this Explicit Mean Equation outperforms the original implicit formulation of Loewe Additivity and Bliss Independence when measuring synergy in terms of the deviance between measured and expected response. Further, we show that a deviance based computation of synergy outperforms a parametric approach. We show this on two datasets of compound combinations that are categorized into synergistic, non-interactive and antagonistic [Yadav et al., 2015, Cokol et al., 2011].

## 1 Introduction

When combining a substance with other substances, one is generally interested in interaction effects. Those interaction effects are usually described as synergistic or antagonistic, dependent on whether the interaction is positive, resulting in greater effects than expected, or negative, resulting in smaller effects than expected. From data generated with high-throughput techniques, one is confronted with massive compound interaction screens. From those screens, one needs to filter for interesting candidates that exhibit an interaction effect. To quickly scan all interactions, a simple measure is needed. Based on that pre-processing scan, those filtered combination candidates can then be examined in greater detail.

To determine whether a combination of substances exhibits an interaction effect, it is crucial to determine a non-interactive effect. Only when deviance from that so-called null reference is observed, can one speak of an interactive effect [Lederer et al., 2018a]. Over the last century, many principles of non-interaction have been introduced. For an extensive overview, refer to [Greco et al., 1995, Geary, 2012]. Two main principles for non-interactivity have survived the critics: Loewe Additivity [Loewe, 1928] and Bliss Independence [Bliss, 1939]. The popularity of Loewe Additivity is based on its principle of sham combination which assumes no interaction when a compound is combined with itself. Other null reference models do not hold that assumption. An alternative is Bliss Independence, which assumes (statistical) independence between the combined compounds.

Independent of the indecisive opinions about the null reference, there are multiple proposals how synergy can be measured given a null reference model. Some suggest to measure synergy as the difference between an observed isobole and a reference isobole calculated from a null-reference model. An isobole is the set of all dose combinations of the compounds that reach the same fixed effect, such as 50% of the maximal effect [Minto et al., 2000, Chou and Talalay, 1984]. Another way to quantify synergy on the basis of the isobole is to look at the curvature and arc-length of the longest isobole spanned over the measured response [Cokol et al., 2011]. As the deviation from an isobole is measured for a fixed effect or dose ratio, synergy is only measured locally along that fixed effect or dose ratio. In order to not miss any effects, this method has to be applied for as many dose ratios possible.

In this paper we measure synergy as the deviation over the entire response surface. One way to do so is the Combenefit method by measuring synergy in terms of volume between the expected and measured effect [Di Veroli et al., 2016]. We will refer to it as a lack-of-fit method as it quantifies the lack of fit from the measured data to the null reference model. Another way of capturing the global variation is by introducing a synergy parameter *α* into the mathematical formulation of the response surface. This parameter *α* is fitted by minimizing the error between the measured effect and the *α*-dependent response surface. Such statistical definition of synergy allows for statistical testing of significance of the synergy parameter. Fitting a synergy parameter to the data as in the parametric approaches tends to be computationally more complex than computing the difference between the raw data and the null model as in the lack-of-fit approaches.

There is an increase in theoretical approaches to synergy, such as the recently re-discovered Hand model [Hand, 2000, Sinzger et al., 2019], which is a formulation of Loewe Additivity in form of a differential expression, or new ways of defining and measuring synergy, such as the ZIP model [Yadav et al., 2015], SynergyFinder [He and Tang, 2016], MuSyC [Meyer et al., 2019] and the copula model [Lambert and Dawson, 2019]. It would be a large effort to compare these recent approaches with ours. An extensive comparison of the models has recently been made in Meyer et al. [2019]. Hence we focus on the two main principles, Loewe Additivity and Bliss Independence.

As the research area of synergy evolved from different disciplines, different terminologies are in common use. Whilst in pharmacology, one refers to the Loewe model, in toxicology, the same principle is called concentration addition. The response can be measured among others in growth rate, survival, or death. It is usually referred to as the measured or phenotype effect or as cell survival. In this study we interchange the terms response and effect.

When measuring a compound combination, one also measures each agent individually. The dose or concentration is typically some biological compound per unit of weight when using animal or plant models or per unit of volume when using a cell-based assay. However, it can also be an agent of a different type for example a dose of radiation as used in modern combination therapies for cancer [Nat, 2018]. This individual response is called mono-therapeutic response [Di Veroli et al., 2016] or single compound effect. We prefer a more statistical terminology and refer to it as conditional response or conditional effect. With record we refer to all measurements taken of one cell line or organism which is exposed to all combinations of two compounds. In other literature, this is referred to as response matrix [Lehar et al., 2007, Yadav et al., 2015].

In Section 2.1, we give a short introduction to the two null response principles, Loewe Additivity and Bliss Independence. We explain in detail several null reference models that build on those principles. We introduce synergy as any effect different from an interaction free model in Section 2.2. There, we also introduce the parametrized and deviance based synergy approaches. In Section 2.3, we introduce two datasets that come with a categorization into synergistic, non-interactive and antagonistic. We evaluate the models and methods in Section 3 together with a detailed comparison of the synergy scores.

## 2 Materials and Methods

### 2.1 Theory

Before one can decide whether a compound combination exhibits a synergistic effect, one needs to decide on the expected effect assuming no interaction between the compounds. Such so-called null reference models are constructed from the conditional (mono-therapeutic) dose-response curves of each of the compounds, which we denote by *f*_*j*_ (*x*_*j*_) for *j* ∈ {1, 2}. Null reference models extend the conditional dose-response curves to a (null-reference) surface spanned between the two conditional responses. We denote the surface as *f* (*x*_1_, *x*_2_) such that

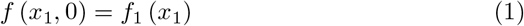

and

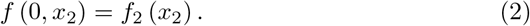

Thus, the conditional response curves are the boundary conditions of the null reference surface. For this study, we focus on Hill curves to model the conditional dose-responses. More detailed information can be found in Appendix A.

#### 2.1.1 Loewe Additivity

Loewe Additivity builds on the concepts of sham combination and dose equivalence. The first concept is the idea that a compound does not interact with itself. The latter concept assumes that both compounds that reach the same effect can be interchanged. Therefore, any linear combination of fractions of those doses which reach the effect individually and, summed up, are equal to one, yields that exact same effect. Mathematically speaking, if dose 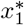 from the first compound reaches the same effect as dose 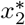 from the second compound, then any dose combination (*x*_1_, *x*_2_), for which

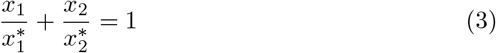

holds, should yield the same effect as 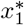 and 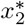. As this idea can be generalized to any effect *y*, one gets

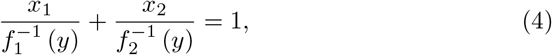

where 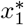 and 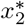 are replaced with 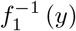 and 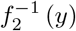, the inverse functions of Hill curves, respectively. For a fixed effect *y*, Eq. 4 defines an isobole, which is in mathematical terms a contour line. Hence the name of this model: the General Isobole Equation. It is an implicit formulation as the effect *y* of a dose combination (*x*_1_, *x*_2_) is implicitly given in Eq. 4. In the following we use the mathematical notation for the General Isobole Equation *f*_GI_ (*x*_1_, *x*_2_) = *y* with *y* being the solution to Eq. 4.

It was shown by Lederer et al. [2018a] that the principle of Loewe Additivity is based on a so-called Loewe Additivity Consistency Condition (LACC). This condition is that it should not matter whether equivalent doses of two compounds are expressed in terms of the first or the second. Under the assumption of the LACC being valid, Lederer et al. [2018a] have shown, that a null reference model can be formulated explicitly, by expressing the doses of one compound in terms of the other compound:

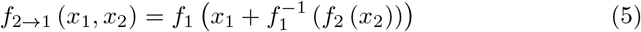

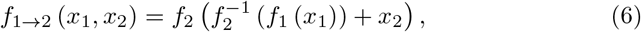

where 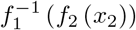 is the dose *x*_1_ of compound one to reach the same effect of compound two with dose *x*_2_ (see Fig. 7 in Appendix A). For a detailed explanation, refer to Lederer et al. [2018a]. Summing up this dose equivalent of the first compound with the dose of the first compound allows for the computation of the expected effect of the compound combination. With the two formulations above, the effect *y* of the dose combination (*x*_1_, *x*_2_) is expressed as the effect of either one compound to reach that same effect. Under the LACC, all three models, Eq. 4, Eq. 5 and Eq. 6 are equivalent. It was further shown, that, in order for the LACC to hold, conditional dose-response curves must be proportional to each other, i.e. being parallel shifted on the *x*-axis in log-space. It has been commented by Geary [2012] and shown in [Lederer et al., 2018a], that this consistency condition is often violated. In an effort to take advantage of the explicit formulation and to counteract the different behavior of Eq. 5 and Eq. 6 in case of a violated LACC, Lederer et al. [2018a] introduced the so-called Explicit Mean Equation as mean of the two explicit formulations of Eq. 5 and Eq. 6:

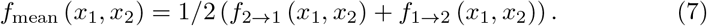

A more extensive overview of Loewe Additivity and definition of null reference models together with visualizations can be found in Lederer et al. [2018a].

#### 2.1.2 Bliss Independence

Bliss Independence assumes independent sites of action of the two compounds and was introduced a decade later than Loewe Additivity in [Bliss, 1939]. Note that the formulation of Bliss Independence depends on the measurement of the effect. The best known formulation of Bliss Independence is based on monotonically increasing responses for increasing doses:

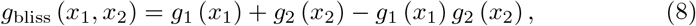

where *g_i_* (*x_i_*) = 1 − *f_i_* (*x_i_*) is a conditional response curve with increasing effect for increasing doses. In case the effect is measured in percent, i.e. *y* ∈ [0, 100], the interaction term needs to be divided by 100 to ensure the right dimensionality of the term.

Here, we measure the effect in terms of cell survival or growth inhibition. Therefore the conditional response curves are monotonically decreasing for increasing concentrations or doses.

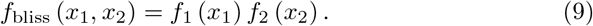

The records are normalized to the response at *x*_1_ = 0, *x*_2_ = 0, thus *f*_1_ (0) = *f*_2_ (0) = 1. To arrive from Eq. 8 to Eq. 9, one replaces any *g* by 1 − *f*. Chou and Talalay [1984] derive the Bliss Independence from a first order Michaelis-Menten kinetic system with mutually non-exclusive inhibitors.

### 2.2 Methods

The models introduced in the previous section are null reference models in that they predict a response surface in the absence of compound interaction. We capture synergy in a single parameter to facilitate the screening process. This is different from other approaches, such as Chou and Talalay [1977], who measure synergy as deviation from a null-reference isobole without summarizing the deviation in a single parameter. The single parameter value is typically referred to as synergy- or *α*-score [Berenbaum, 1977]. As we investigate two methods to quantify synergy, we introduce two synergy parameters *α* and *γ*, which measure the extent of synergy. Both synergy scores *α* and *γ* are parametrized such that *α* = 0 or *γ* = 0 denote absence of an interaction effect. In case *α* or *γ* take a value different from zero, we speak of a non-additive, or interactive effect. A compound combination is, dependent on the sign of synergy parameter, one of the three following:

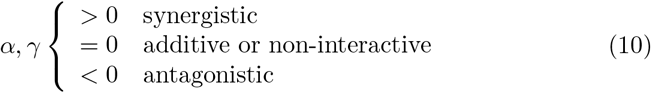

Here, we measure synergy in two different ways, namely in fitting parametrized models or computing the lack-of-fit. The first method fits null reference models that are extended with a synergy parameter *α*. For these parametrized models *α* is computed by minimizing the square deviation between the measured response and the response spanned by the *α*-dependent model. For the second method the difference between a null reference model and the data is computed. For this method, the synergy score *γ* is defined as the volume that is spanned between the null reference model and the measured response.

Just as the conditional responses form the boundary condition for the null-reference surface (Eq. 1, Eq. 2), we want the conditional responses to be the boundary condition for all values of *α*. Explicitly, assuming a synergy model dependent on *α* is denoted by *f* (*x*_1_, *x*_2_|*α*), then

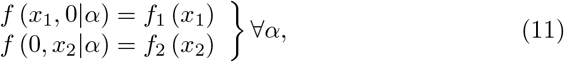

with *f_i_* denoting the conditional response of compound *i*. We refer to Eq. 11 as the Synergy Desideratum. As we will see below, not all synergy models fulfill this property.

#### 2.2.1 Parametrized Synergy

We extend the null reference models introduced in Section 2.1 in Eq. 4 - Eq. 9 to parametrized synergy models. The extension of the General Isobole Equation is the popular Combination Index introduced by Berenbaum [1977] and Chou and Talalay [1984]:

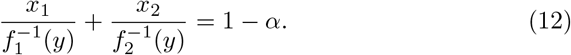

Berenbaum originally equated the left-hand side of Eq. 4 to the so-called Combination Index *I*. Depending on *I* smaller, larger, or equal to 1, synergy, antagonism or non-interaction is indicated. For consistency with the other synergy models, we set *I* = 1 − *α* such that *α* matches the outcomes as listed in Eq. 10. In Section 3 we will refer to this implicit model as *f*_CI_ (*x*_1_, *x*_2_|*α*), where *α* is the parameter that minimizes the squared error between measured data and Eq. 12.

Note that this model violates the Synergy Desideratum in Eq. 11 as *α* not zero leads to deviations from the conditional responses. Explicitly, *f*_CI_ (*x*_1_, 0|*α*) = *f*_1_ ((1 − *α*) *x*_1_) ≠ *f*_1_ (*x*_1_). Although the Combination Index model violates the Synergy Desideratum, in practice it performs quite well and is in widespread use.

The explicit formulations in Eq. 5 and Eq. 6 are equivalent to the General Isobole Equation, *f*_GI_ (*x*_1_, *x*_2_), given in Eq. 4, under the LACC [Lederer et al., 2018a], but different if the conditional responses are not proportional. The two explicit equations are in fact an extension of the ‘cooperative effect synergy’ proposed by Geary [2012] for compounds with qualitatively similar effects. For these explicit formulations in Eq. 5 and Eq. 6 we propose a model that captures the interaction based on the explicit formulations:

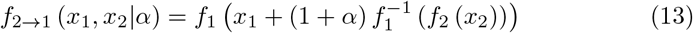

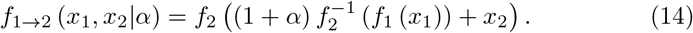

With this, we can extend the Explicit Mean Equation model *f*_mean_ (*x*_1_, *x*_2_) in Eq. 7 to a parametrized synergy model:

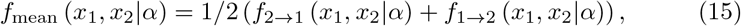

which we refer to as *f*_mean_ (*x*_1_, *x*_2_|*α*). As *f*_2→1_ (*x*_1_, *x*_2_|*α*) and *f*_1→2_ (*x*_1_, *x*_2_|*α*) do not fulfill the Synergy Desideratum, *f*_mean_ (*x*_1_, *x*_2_|*α*) also does not fulfill it.

To investigate the difference between the two models *f*_2→1_ (*x*_1_, *x*_2_) (Eq. 5) and *f*_1→2_ (*x*_1_, *x*_2_) (Eq. 6) we treat compound one and two based on the difference in slopes in the conditional responses (for more detailed information on the different parameters in Hill curves, refer to Appendix A). Instead of speaking of the first and second compound, we speak of the smaller and larger one, referring to the order of steepness. Therefore, we use models Eq. 13 and Eq. 14, but categorize the compounds based on the slope parameter of their conditional response curves. This results in *f*_large→small_ (*x*_1_, *x*_2_|*α*) and *f*_small→large_ (*x*_1_, *x*_2_|*α*).

Another synergy model we introduce here and refer to as *f*_geary_ (*x*_1_, *x*_2_ *α*) is based on a comment of Geary [2012], hence the naming. The two explicit models *f*_2→1_ (*x*_1_, *x*_2_) and *f*_1→2_ (*x*_1_, *x*_2_) yield the same surface under the LACC but do rarely in practice. Therefore, it cannot be determined whether a response that lies between the two surfaces is synergistic or antagonistic and hence should be treated as non-interactive. Thus, if *α* from *f*_1→2_ (*x*_1_, *x*_2_|*α*) and *α* from *f*_2→1_ (*x*_1_, *x*_2_ *α*) are of equal sign, the synergy score of that model is computed as the mean of those two parameters. In case the two synergy parameters are of opposite sign, the synergy score is set to 0:

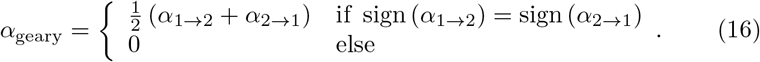

Next, to extend the null reference model following the principle of Bliss Independence, we extend Eq. 8 to

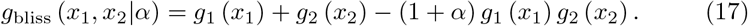

The motivation for this model is that any interaction between the two compounds is caught in the interaction term of the two conditional responses. In case of no interaction, the synergy parameter *α* = 0, which leads to (1 + *α*) = 1, and results in no deviance from the null reference model. As we use the formulation of Eq. 9 due to measuring the effect as survival, we reformulate Eq. 17 analogously as we did to get from Eq. 8 to Eq. 9: by replacing *g_i_* (*x_i_*) with 1 − *f_i_* (*x_i_*). Hence, Eq. 17 takes the form:

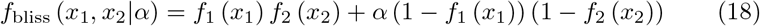

This model does satisfy the requirement of no influence of the synergy parameter on conditional doses: *f*_bliss_ (*x*_1_, 0|*α*) = *f*_1_ (*x*_1_) and *f*_bliss_ (0, *x*_2_|*α*) = *f*_2_ (*x*_2_) as *f_i_* (0) = 1. In case of synergy, the interactive effect is expected to be larger, therefore, *α* being positive. If the compound combination has an antagonistic effect, the interaction term is expected to be negative. For extreme *α*, the parametric approach leads to responses outside of the range 0 ≤ *y* ≤ 1, e.g. *f*_bliss_ (*x*_1_, *x*_2_) → −∞ if *α* → −∞. The same holds for the formulations of Loewe Additivity. The implicit formulation becomes impossible to match and for the explicit formulations, the dose expression within brackets of *f*_2→1_ (*x*_1_, *x*_2_|*α*) becomes negative. Additionally, *α* > 1 is not possible for *f*_CI_ (*x*_1_, *x*_2_|*α*), as the left-hand side of Eq. 12 can not be negative. Such behavior is also known from other models, e.g. for the Greco flagship model for negative synergy scores [Greco et al., 1995, p. 365-366, and Fig. 26]. Hence, we will limit *α* to the range of −1 to 1.

Despite of the Synergy Desideratum being violated for the models that build up on the Loewe Additivity principle, there is no further effect on the model comparison presented in Section 3 as conditional doses are excluded when computing the synergy score (see Section 2.2.2 and Section 2.2.3).

#### 2.2.2 Lack-of-Fit Synergy

The second method to measure synergy investigated here is to compute the lack-of-fit of the measured response of a combination of compounds to the response of a null reference model derived from the conditional responses. We refer to this synergy value as *γ*:

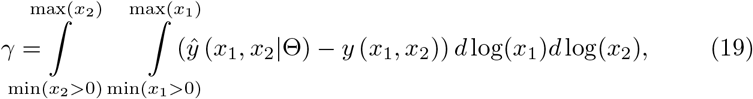

with 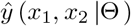 the estimated effect with parameters Θ of the fitted conditional responses following any non-interactive model and *y* the measured effect. Note that 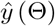 and *y* are dependent on the concentration combination (*x*_1_, *x*_2_). This method was used in the AstraZeneca DREAM challenge [Menden et al., 2018] with the General Isobole Equation as null reference model and can be found in [Di Veroli et al., 2016]. Computing the volume has the advantage of taking the experimental design into account in contrast to simply taking the mean deviance over all measurement points, which is independent of the relative positions of the measurements. We also used a synergy value calculated from the mean deviance and it clearly performed worse (data not shown). The synergy value varies for different dose transformations. For example, the computed null-reference surface (and hence the synergy value) will be different for the same experiment if a log-transformation is applied to the doses or not.

In all, we have introduced six null reference models, five of them building up on the concept of Loewe Additivity and one on Bliss Independence. We further have introduced two methods to compute synergy, the parametric one and the lack-of-fit method, where both synergy parameters *α* and *γ* are positive if the record is synergistic, negative, if antagonistic. This results in twelve synergy model-method combinations: the parametric ones, *f*_CI_ (*x*_1_, *x*_2_|*α*) (Eq. 12), *f*_large→small_ (*x*_1_, *x*_2_|*α*) and *f*_small→large_ (*x*_1_, *x*_2_|*α*) (Eq. 13, Eq. 14, dependent on the slope parameters) together with their mean, *f*_mean_ (*x*_1_, *x*_2_|*α*) (Eq. 15), the method of Geary and *f*_bliss_ (*x*_1_, *x*_2_|*α*) (Eq. 17). For the lack-of-fit method, we take as the null reference: *f*_GI_ (*x*_1_, *x*_2_) (Eq. 4), *f*_large→small_ (*x*_1_, *x*_2_) and *f*_small→large_ (*x*_1_, *x*_2_) (Eq. 5, Eq. 6), with the Explicit Mean Equation, *f*_mean_ (*x*_1_, *x*_2_) (Eq. 7), the method of Geary (analogously to Eq. 16) and *f*_bliss_ (*x*_1_, *x*_2_) (Eq. 9).

#### 2.2.3 Fitting the Synergy Parameter

Before applying the two methods presented in Section 2.2.1 and Section 2.2.2, we normalize and clean the data from outliers. In a first step we normalize all records to the same value, *y*_0_, the measured response at zero dose concentration from both compounds. Second, we discard outliers using the deviation from a spline approximation. Third, we fit both conditional responses of each record, namely the responses of each compound individually, to a pair of Hill curves (Eq. 21, Appendix A). We fit the response at zero dose concentration for both Hill curves. This gives the parameter set **Θ** = *y*_0_, *y_∞,_*_1_, *y_∞,_*_2_, *e*_1_, *e*_2_, *s*_1_, *s*_2_ for each record. More details are given in Appendix B.

We apply the two different methods to calculate the synergy parameters *α* and *γ* to each record. First, for the parametrized synergy models, we apply a grid search for *α*, for *α* ∈ [−1, 1] with a step size of 0.01, minimizing the sum of squared errors. This gives the value of *α* for which the squared error between the *i*^th^ measured effect *y*^(*i*)^ and *i*^th^ expected effect 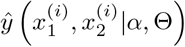 is minimal:

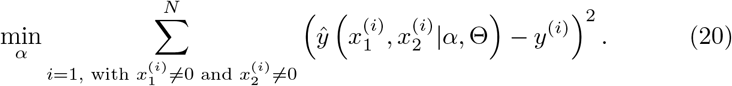

Note that we exclude the conditional responses that we used to fit **Θ** from the minimization. Second, we apply the lack-of-fit method from Di Veroli et al. [2016], where synergy is measured in terms of the integral difference in log space of measured response and surface spanned by the non-interactive models in Section 2.1, as given in Eq. 19. For the calculation of the integrals, we apply the trapezoidal rule [Press et al., 2007, Chapter 4]. In Fig. 1 we summarize the most important steps of the analysis for a synergistic example. In Appendix E, Fig. 11, the same is shown for an antagonistic record.

**Figure 1:**
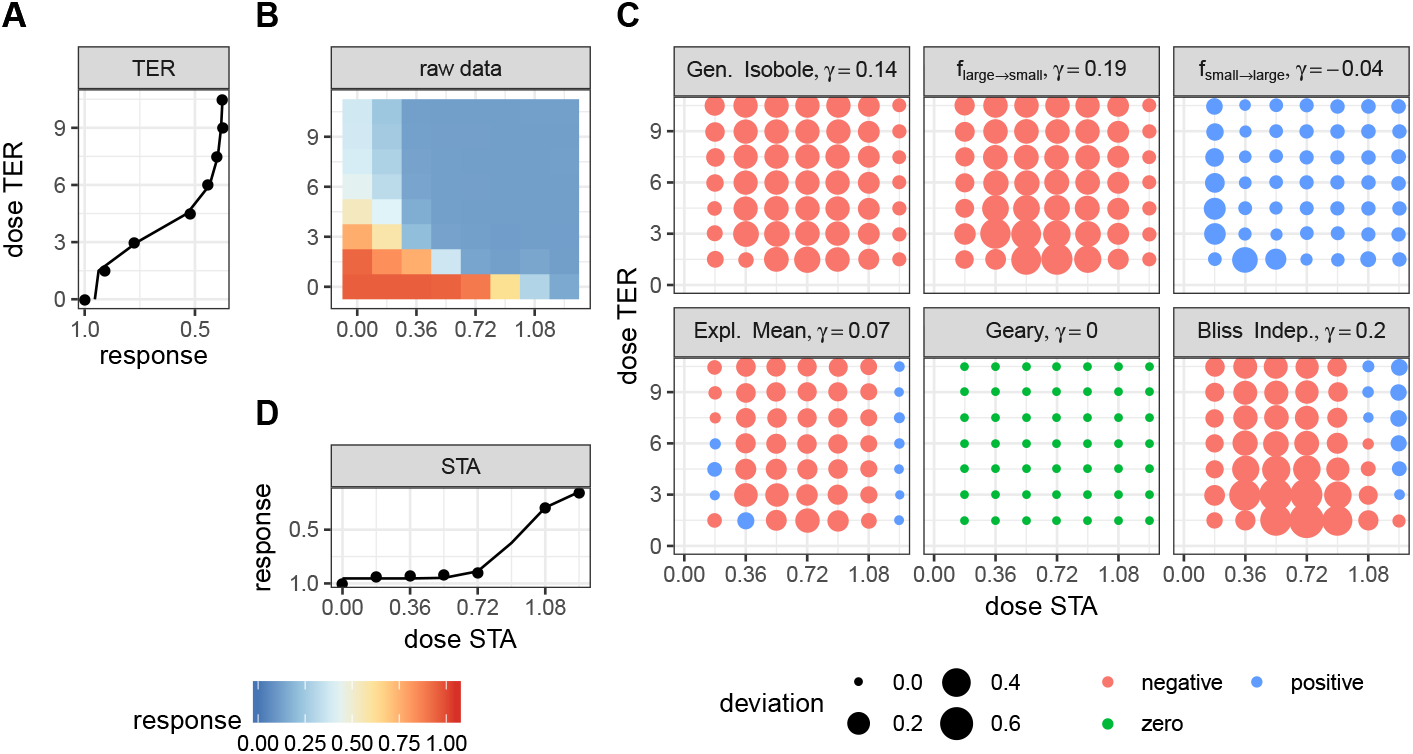
Description of the analysis steps of the lack-of-fit method for the compound pair TER and STA from the Cokol dataset. This compound pair is categorized as synergistic according to [Cokol et al., 2011]. The raw response data of the record is depicted in (B). The response data normalized by the read at zero dose concentration (lower left). In (B) the degree of relative cell growth is colored from high to low values in red to blue. Step 1: compute Hill curves for conditional responses: Based on the raw reads of the single dose responses (lower and left outer edges) fit a Hill curve to the conditional responses. The fitted Hill curves shown in (A) and (D) with original raw data shown as points. Step 2: compute expected non-interactive response for all six models: not shown. Step 3: compute difference between measured data (C) and expected data from all six null reference models: shown in (C). The direction of difference is shown by color (red for negative and blue for positive, green for zero). The larger the degree of difference, the larger the bullet, and vice versa. Step 4: compute integral *γ* over the differences: Over all those bullets, we then compute the integral, which gives the synergy score *γ*. For every model, the synergy score *γ* is depicted in the title of each matrix in (C).

### 2.3 Material

To evaluate the two methods introduced in Section 2.2.1 and Section 2.2.2, we apply them to two datasets of compound combination screening for which a categorization into the three synergy cases is provided.

The Mathews Griner dataset is a cancer compound synergy study by Mathews Griner et al. [2014]. In a one-to-all experimental design, the compound ibrutinib was combined with 463 other compounds and administered to the cancer cell line TMD8 of which cell viability was measured. The dataset is published at https://tripod.nih.gov/matrix-client/. Each compound combination was measured for 5 different doses, decreasing from 125*μM* to 2.5*μM* in a four-fold dilution for each compound alongside their conditional effects, resulting in 36 different dose combinations. The categorization of this dataset comes from a study by Yadav et al. [2015], in which every record was categorized based on a visual inspection.

The Cokol dataset comes from a study about fungal cell growth of the yeast S. cerevisiae (strain By4741), where Cokol et al. [2011] categorized the dataset. In this study the influence on cell growth was measured when exposed to 33 different compounds that were combined with one another based on promising combinations chosen by the authors, resulting in 200 different drug-drug-cell combinations. With an individually measured maximal effect dose for every compound, the doses administered decrease linearly in seven steps with the eight dose set to zero, resulting in an 8 × 8 factorial design.

Based on the longest arc length of an isobole that is compared to the expected longest linear isobole in a non-interactive scenario, where Loewe Additivity serves as null reference model, each record was given a score. In more detail, from the estimated surface of a record assuming no interaction, the longest contour line is measured in terms of its length and direction (convex or concave). A convex contour line leads to the categorization of a record as synergistic and the arc length of the longest contour line determines the strength of synergy. A concave contour line results in an antagonistic categorization with its extent being measured again as the length of the longest isobole. Thus the Cokol dataset not only comes with a classification but also with a synergy score similar to *α* or *γ*.

To our knowledge, these two datasets are the only high-throughput ones with a classification into the three synergy classes: antagonistic, non-interactive and synergistic. Both datasets are somewhat imbalanced because interactions are rare [Borisy et al., 2003, Zhang et al., 2007, Farha and Brown, 2010]. The distribution of the classification is listed in Table 1. We obtained both categorizations after personal communication with the authors Yadav et al. [2015] and Cokol et al. [2011]. For the purpose of comparing the synergy models, we consider these two classifications as ground truth.

**Table 1:** Number of cases categorized as synergistic, antagonistic or non-interactive in the two datasets Mathews Griner and Cokol.

|  | synergistic | antagonistic | non-interactive |
| --- | --- | --- | --- |
| Mathews Griner | 121 | 90 | 252 |
| Cokol | 50 | 68 | 82 |

## 3 Results

Using the two methods of computing the synergy score, the parametric one (Section 2.2.1) and the lack-of-fit one (Section 2.2.2), we compute synergy scores for all records of the two datasets introduced in Section 2.3.

### 3.1 Kendall rank correlation coefficient

Having computed the synergy scores *α* and *γ* from the two different methods as described in Section 2.2.3, we compute the Kendall rank correlation coefficient, which is also known as Kendall’s tau coefficient and was originally proposed by Kendall [1938]. This coefficient computes the rank correlation between the data as originally categorized by Yadav et al. [2015] and Cokol et al. [2011] and the computed synergy scores resulting from the two methods introduced in Section 2.2.1 and Section 2.2.2. For the analysis, we rank synergistic records highest at rank 3, followed by non-interactive at rank 2 and antagonistic lowest at rank 1. Due to the many ties in rank, the Kendall rank correlation coefficient cannot take a value higher than 0.75 for Mathews Griner and 0.8 for Cokol, even if a perfect ranking was given. An overview of the Kendall rank correlation coefficients is given in Table 3 and Table 4 in Appendix D.

To compare the parametric and lack-of-fit methods, we plot the correlation values as a scatter plot per method (see Fig. 2) with the values from the parametric method plotted on the *x*-axis and those from the lack-of-fit method on the *y*-axis. Most of the points scatter in the upper left triangle, above the diagonal line. This shows that the lack-of-fit method outperforms the parametric method. This holds for all models applied to the Mathews Griner dataset and also for all models but *f*_geary_ (*x*_1_, *x*_2_|*α*) and *f*_small→large_ (*x*_1_, *x*_2_|*α*) applied to the Cokol dataset. For both datasets, the highest correlation scores result from those null reference models that are based on the Loewe Additivity principle. The Bliss null reference model performs worst for the Mathews Griner set for both methods. For the Cokol data it is the second worst model. To a certain extent this can be explained due to the classification of the Cokol dataset being based on the isobole length relative to non-interactive isoboles, which is a Loewe Additivity type analysis. As the categorization of the Mathews Griner dataset is based on visual inspection, we cannot explain the bad performance of *f*_bliss_ (*x*_1_, *x*_2_) for that dataset. On both datasets, *f*_GI_ (*x*_1_, *x*_2_), *f*_large→small_ (*x*_1_, *x*_2_) and *f*_mean_ (*x*_1_, *x*_2_) perform best for the lack-of-fit method. For the Mathews Griner dataset, *f*_large→small_ (*x*_1_, *x*_2_) dominates marginally over the General Isobole Equation and Explicit Mean Equation model. For the Cokol dataset, the Explicit Mean Equation dominates for both methods.

**Figure 2:**
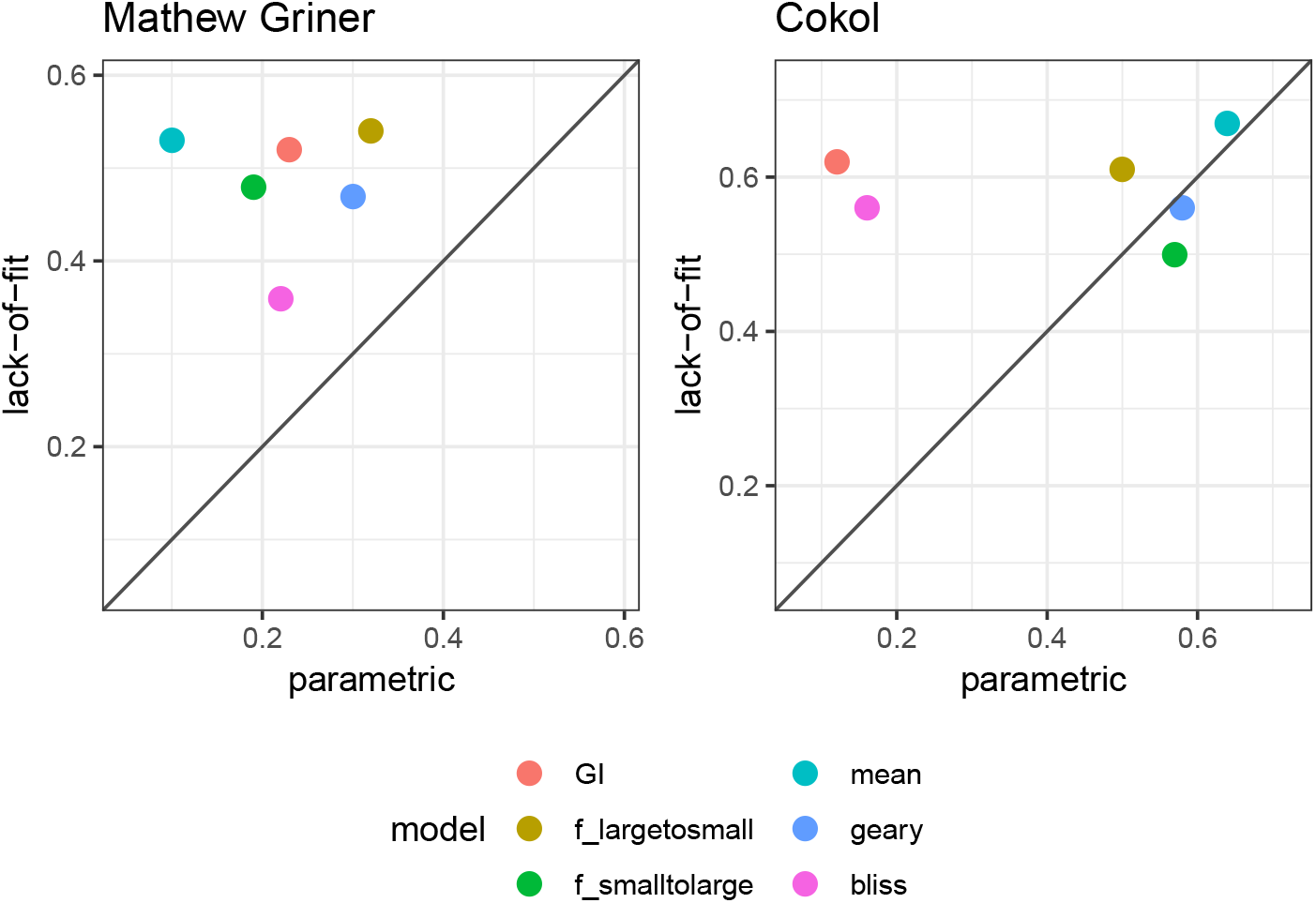
Scatter plot of Kendall rank correlation coefficient for both datasets, Mathews Griner (left) and Cokol (right). The Kendall correlation measures the rank correlation of the original categorization and the computed synergy scores. The higher the correlation, the more similar the score ranking. The correlation values from the synergy scores *α*, computed with the parametric approach, are plotted on the *x*-axis and those from the lack-of-fit approach are plotted on the *y*-axis. Each model is depicted in a different color. To guide the eye, the diagonal is plotted. If a data point is above the diagonal, the Kendall rank correlation coefficient from the lack-of-fit method is higher than that from the parametric method, and vice versa. Without exception, the Kendall rank correlation coefficients are all higher for the synergy scores *γ*, which are computed with the lack-of-fit method, than those based on the *α* scores computed with the parametric method.

### 3.2 Scattering of Synergy Scores

To further investigate the performance of the methods and null reference models, we plot the synergy scores of the best performing models based on the Kendall rank correlation coefficient analysis (Section 3.1, and an ROC analysis, which we describe in detail in Appendix C) for both datasets in Fig. 3, Fig. 4 and Fig. 5. In all figures, the overall correlation of the compared data is depicted together with the correlation per categorization. The coloring of the scores is based on the original categorization as antagonistic, non-interactive or synergistic as provided by Yadav et al. [2015] and Cokol et al. [2011].

**Figure 3:**
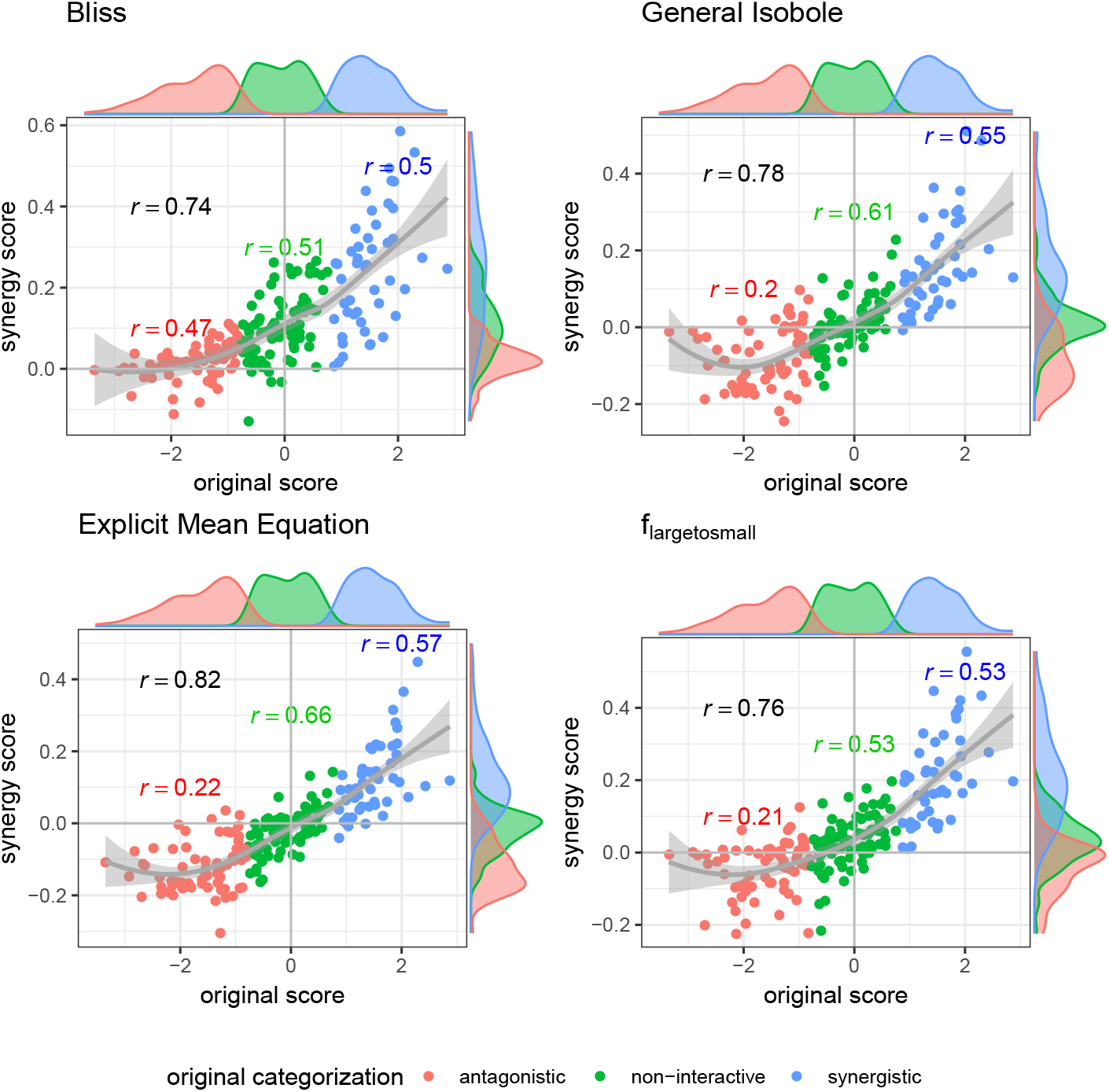
Computed synergy scores *γ* of Cokol data of the best models according to the Kendall rank correlation coefficient and ROC analysis in Section 3.1 and Appendix C in comparison to the original scores from Cokol et al. [2011]. The data points are colored based on the original categorization. For all three categories, synergistic, non-interactive and antagonistic, the Pearson correlation is depicted between the original scores in that category and the computed synergy scores in the respective color. Additionally, we depict the local polynomial regression fitting of all scores (in gray). The histograms of the scores are plotted on the axis, separated by color based on the original categorization. Synergy scores *γ* based on the Explicit Mean Equation model show the highest correlation with the original scores.

**Figure 4:**
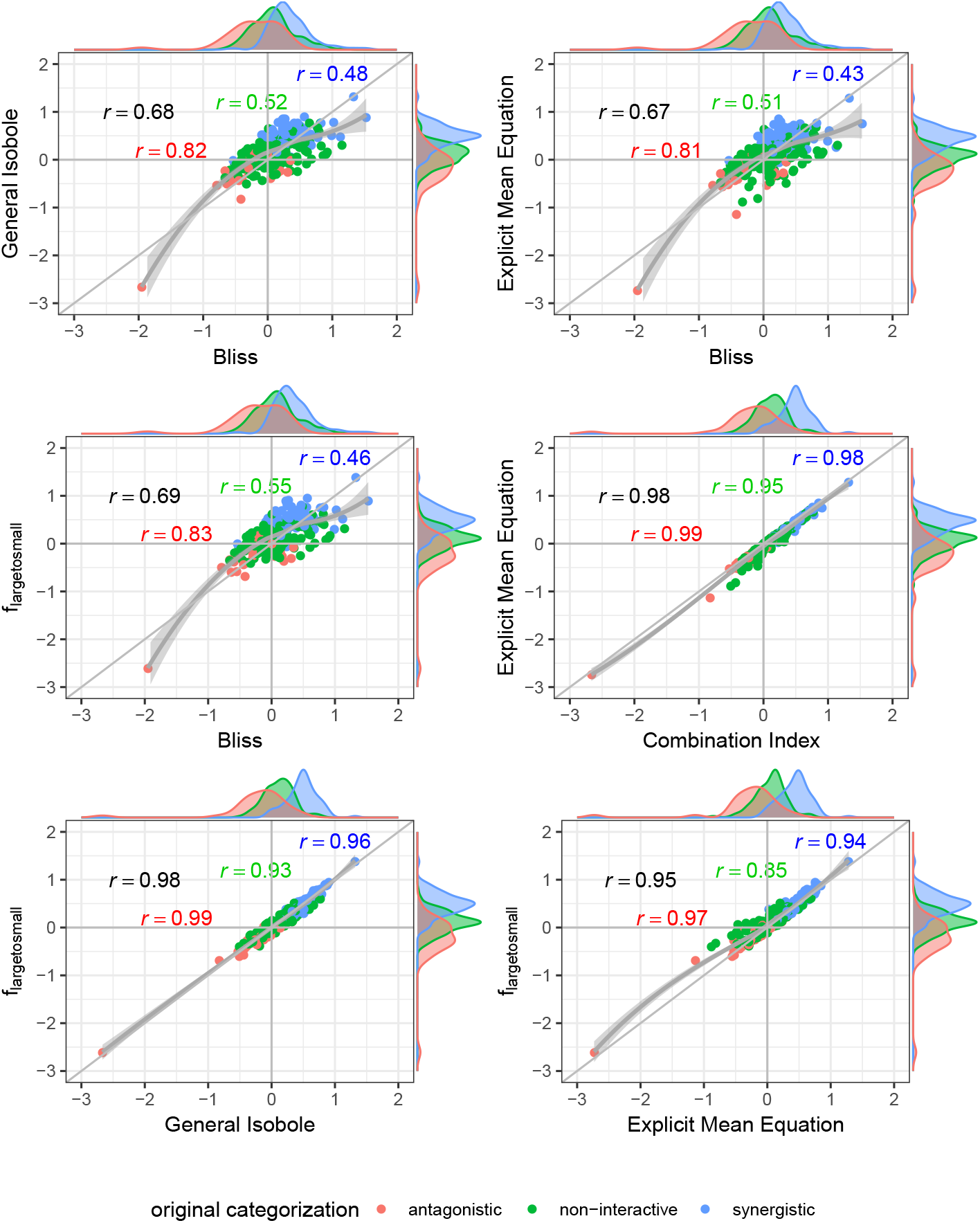
Scatter plot of synergy scores *γ* of the Mathews Griner dataset. The scores are computed with the lack-of-fit method. Displayed are the four best models according to the Kendall rank correlation coefficient and ROC analysis in Section 3.1 and Appendix C. The scores of one model are depicted on the *x*-axis and the other on the *y*-axis. The original categorization is given based on colour. The Pearson correlation score between the synergy scores are depicted by color for every categorization and the overall Pearson correlation is depicted in black. To guide the eye, the axis at 0, the diagonal and local polynomial regression fitting are depicted in grey. The histograms of the scores are plotted on the axis, separated by color based on the original categorization. The three models based on the Loewe Additivity principle show highest correlation (center right and lower row). All comparison with *f*_bliss_ (*x*_1_, *x*_2_) show lowest correlation (first three cases). There is a large difference between the correlation between the additive models and the comparison of Bliss Independence by roughly 0.3.

**Figure 5:**
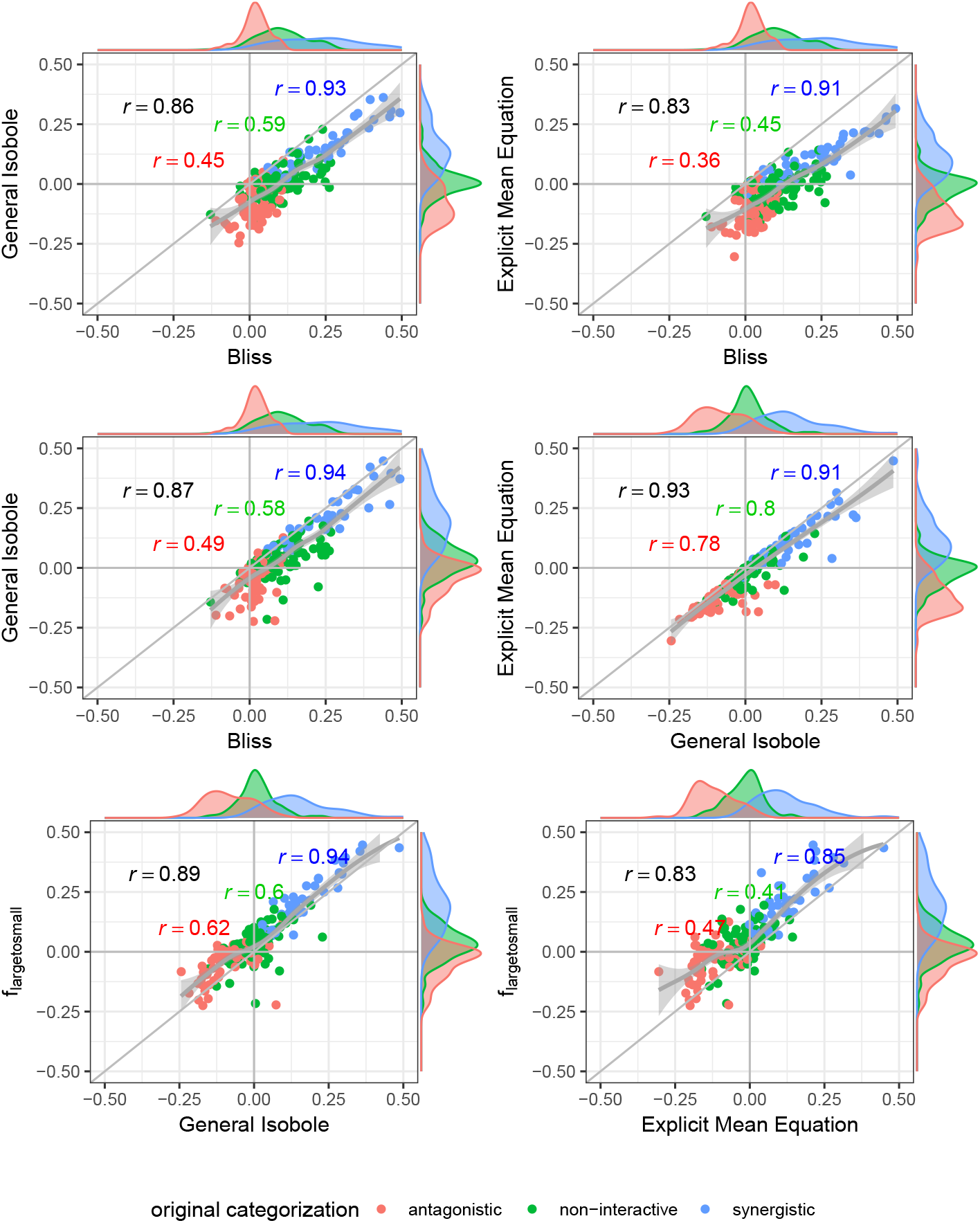
Scatter plot of synergy scores *γ* of the Cokol dataset. The scores are computed with the lack-of-fit method. Displayed are the four best models according to the Kendall rank correlation coefficient and ROC analysis in Section 3.1 and Appendix C. The scores of one model are depicted on the *x*-axis and the other on the *y*-axis. The original categorization is given based on colour. The Pearson correlation score between the synergy scores are depicted by color for every categorization and the overall Pearson correlation is depicted in black. To guide the eye, the axis at 0, the diagonal and the local polynomial regression fitting are depicted in grey. The histograms of the scores are plotted on the axis, separated by color based on the original categorization. *f*_mean_ (*x*_1_, *x*_2_) and *f*_GI_ (*x*_1_, *x*_2_) show highest correlation (center right), *f*_bliss_ (*x*_1_, *x*_2_) shows lowest (first three comparison cases).

In Fig. 3 the synergy scores computed with the lack-of-fit method are plotted against the original synergy scores from Cokol et al. [2011]. Applying the lack-of-fit method to the Bliss Independence model (Eq. 9) results in scores which are mainly above zero (Fig. 3, upper left). Further, it can be seen in the density plots along the *y*-axis in Fig. 3, upper left panel, and on the *x*-axis of Fig. 4, both panels in the first row and left panel in the middle row, that the synergy scores that are computed based on the principle of Bliss Independence cannot be easily separated by categorization, making it difficult to come up with a threshold to categorize a record into one of the three synergy categories (synergy, antagonism, non-interaction) given a synergy score.

For the other three models depicted in Fig. 3, that are based on the principle of Loewe Additivity, the synergy scores are more clearly separated. The computed scores of the synergistic records distribute nicely above zero in the upper right corner (categorized as synergistic and computed synergy scores above zero) as well as they scatter in the lower left corner for antagonistic cases. In all those three panels in Fig. 3 we see for the non-interactive records that the computed scores of those three models are both positive and negative ranging roughly between −0.1 and 0.1 symmetrically. Barely any of the computed synergy scores for antagonistic cases are positive. Therefore, the chances of a record being antagonistic if the synergy score is above zero are quite low as well as the risk of categorizing a record as antagonistic if it is synergistic.

We further looked in detail into dose combinations for which both the *f*_GI_ (*x*_1_, *x*_2_ and *f*_mean_ (*x*_1_, *x*_2_) yield positive synergy values for antagonistic cases and into dose combinations for which the *f*_mean_ (*x*_1_, *x*_2_) model results in negative synergy values for records which are labeled as synergistic. In total we found four dose combinations. A visualization of the observed and expected responses based on the Explicit Mean Equation model is shown in Appendix E, Fig. 12. One of them is a compound combined with itself. Hence, per definition of the Loewe Additivity, no interaction is expected. From Fig. 12, one can see why this record was mis-categorized: for high dose combinations, a greater effect is found, which is not found for the conditional runs. Probably, the dose ranges are too small to show such effects. We looked at the conditional responses of the other three dose combinations and observed that for the originally antagonistic records (three out of four) one of the conditional responses exhibits small effects with the maximal response *y_∞_* being above 0.65 (comp. right panel of Fig. 6). That leads to the computed null-reference surface to be quite high and hence causes synergistic scores if any effects are measured that are smaller than 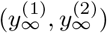. We suspect that the dose concentrations are not well-sampled and larger maximal doses should have been administered.

**Figure 6:**
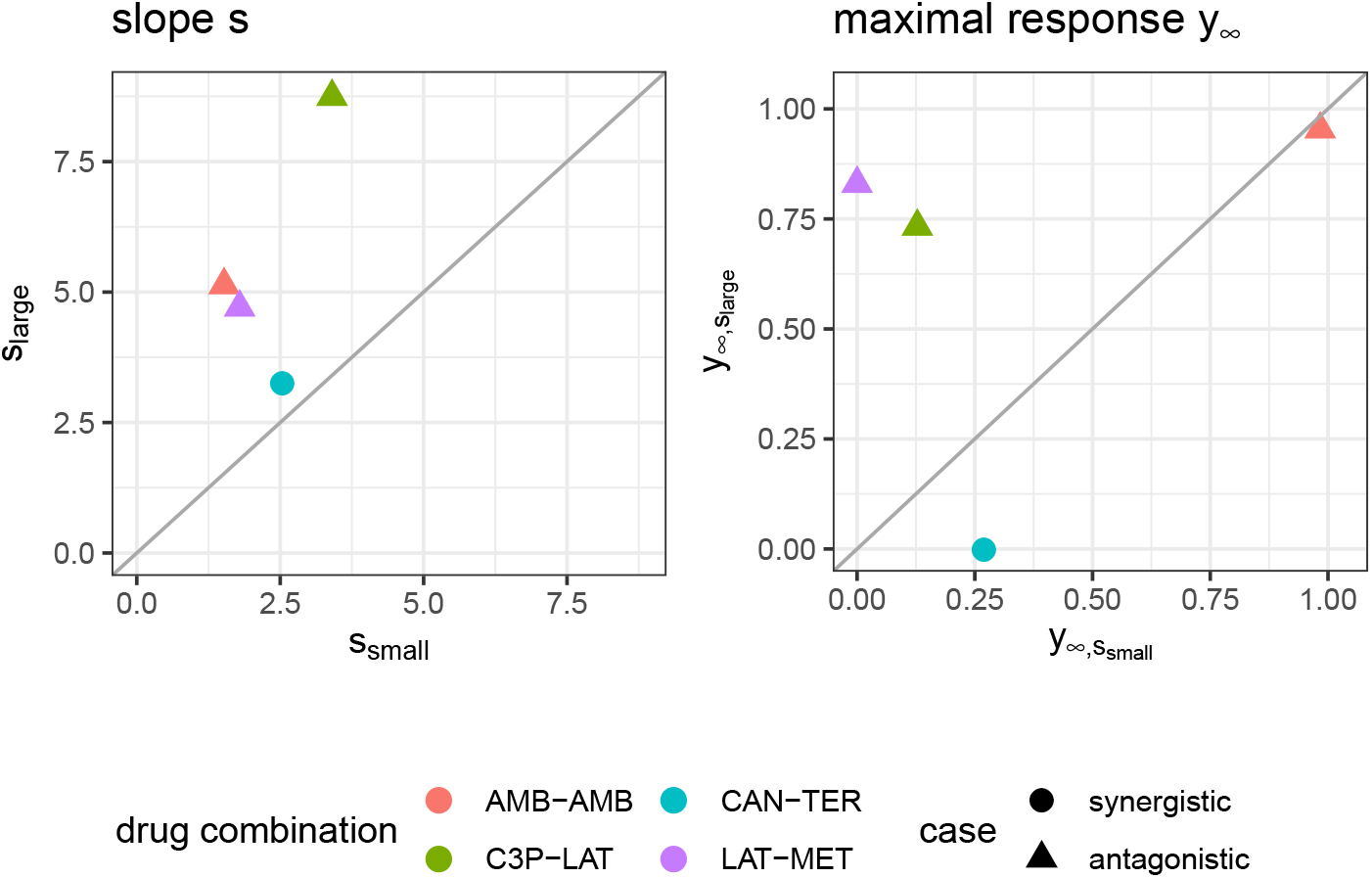
Maximal response *y_∞_* (left) and slope parameters *s* (right) of Hill curves. Parameters are shown for the conditional responses of the four cases for which the lack-of-fit method resulted for *f*_mean_ (*x*_1_, *x*_2_) and *f*_GI_ (*x*_1_, *x*_2_) in a synergy score of opposite sign to its categorization from the Cokol dataset. Different records are depicted in different colours. The original categorization of each record is depicted per shape. The conditional responses of one record, and hence their Hill curve parameters, are grouped depending on size of the Hill curve parameter *s* (larger or smaller).

We further looked up the three dose combinations (excluding the one where the compound is combined with itself) in the Connectivity Map [Subramanian et al., 2017, Lamb et al., 2006], which is one of the largest repositories of drug response studies. Of those, we could find three in the Connectivity Map. All of these dose combinations showed non-interactive effects on all cell lines they were tested on. The assays found in the Connectivity Map are run on cancer cell lines. The dose combinations investigated here are run on yeast. Hence, a full comparison cannot be made, but results are certainly suggestive that the compound combinations are non-interactive.

In Fig. 4 and Fig. 5, the computed scores from different null reference models are plotted against each other. We compare the implicit formulation (General Isobole Equation) to the Bliss Independence model and the two best performing models that are based on the explicit formulation of Loewe Additivity, *f*_mean_ (*x*_1_, *x*_2_) and *f*_large→small_ (*x*_1_, *x*_2_). The coloring of the scores is based on the original categorization as antagonistic, non-interactive or synergistic as provided by Yadav et al. [2015] and Cokol et al. [2011].

In Fig. 4 the scores from the Mathews Griner dataset are plotted. In the two panels in the upper row and the left panel in the middle row Bliss Independence is compared to the other three null models that build up on the principles of Loewe Additivity. It is obvious, that the scores based on Bliss Independence are larger than those of Loewe Additivity and mainly above zero. This is due to the more conservative null reference surface as derived from Bliss Independence (see [Sinzger et al., 2019, Fig. 6]). The scores from models that are based on Loewe Additivity are very similar to each other, as they scatter along the diagonal (panels in middle right and lower row). It is difficult, though, to tell apart whether a record is synergistic or antagonistic, as non-interactive records scatter largely between −0.5 and 0.5. Only records with a computed score outside that range can be categorized as interactive. For the Cokol dataset, which serves as basis for Fig. 5, the scores can be better separated. Despite the scores being generally smaller than those from the Mathews Griner data, the records can be easier separated, when using a Loewe Additivity based model. Additionally, we see here the similarity between these additive models given their strong correlation (right panels in middle row and both panels in lower row). Further, the scores based on *f*_large→small_ (*x*_1_, *x*_2_) achieve higher values than those from the other two Loewe Additivity based models. This becomes obvious when comparing the null-reference surfaces of those three models, as depicted in [Lederer et al., 2018a, Fig. 4]. The surface spanned by *f*_large→small_ (*x*_1_, *x*_2_) spans a surface above those surfaces spanned by Explicit Mean Equation or General Isobole Equation. Therefore, in synergistic cases where the measured effect is greater, and hence the response in cell survival smaller, the difference from the null-reference surface to *f*_large→small_ (*x*_1_, *x*_2_) is greater than to the other two models. We suspect the synergy models from the Cokol dataset to be better separable due to the experimental design of the dataset. All compounds were applied up to their known maximal effect dose. This was not the case for the Mathews Griner dataset, where all compounds were applied at the same fixed dose range.

All in all, the lack-of-fit method performs better for any model when applied to the Mathews Griner dataset and mostly better for the Cokol dataset, with the exception of the *f*_small→large_ (*x*_1_, *x*_2_) and Geary model. We suggest, that the lack-of-fit should be preferred over the parametric method, due to the overall performance on both datasets. When using the lack-of-fit method, the Explicit Mean Equation model performs either second best (Mathews Griner dataset), or best (Cokol dataset). The other two well performing models, the explicit *f*_large→small_ (*x*_1_, *x*_2_) or the original implicit formulation of Loewe Additivity, the General Isobole Equation, do not perform equally well on both datasets. To exclude any bias from these models for different datasets, the Explicit Mean Equation should be preferred.

## 4 Discussion

The rise of high-throughput methods in recent years allows for massive screening of compound combinations. With the increase of data, there is an urge to develop methods that allow for reliable filtering of promising combinations. Additionally, the recent success of a synergy study of in vivo mice by Grüner et al. [2016] underlines the fast development of possibilities to generate biological data. Therefore, it is all the more important to develop methods that are sound and easily applicable to high-throughput data.

In this study we use two datasets of compound combinations that come with a categorization into synergistic, non-interactive or antagonistic for each record.

Based on the fitted conditional responses, we compute the synergy scores of all records. We compare six models that build on the principles of Loewe Additivity and Bliss Independence. Those six models are used with two different methods to compute a synergy score for each record. The first method is a parametric approach and is motivated by the Combination Index introduced by Berenbaum [1977]. The second method quantifies the difference in volume between the expected response assuming no interaction and the measured response and is motivated by Di Veroli et al. [2016].

We compare the computed synergy scores from both methods, each applied with the six reference models, based on Kendall rank correlation coefficients. Based on these correlation coefficients we investigate the reconstruction of ranking of the records (see Section 3.1). We further conduct an ROC analysis (results shown in Appendix C). With this, we quantify the methods’ and models’ capacity to distinguish records from different categories, given a computed synergy score. Both, the Kendall rank correlation coefficient and the ROC analysis, show a superiority of those models that are based on Loewe Additivity relative to those based on Bliss Independence. From those additive models the Explicit Mean Equation is the overall best performing model for both datasets.

For the above comparison of the six null reference models and the two methods, we rely on the underlying categorization of both datasets. All performance metrics are based on how well the predicted synergy scores agree with the underlying categorization. The categorizations of both datasets were created very differently from one another. On one hand, the Mathews Griner dataset was categorized on a visual inspection, on account of which we cannot be certain about the assumptions made that guided the decision making process. On the other hand, the categorization of the Cokol dataset is based on the principle of Loewe Additivity. This leads to the natural preference of null models that are based on Loewe Additivity over those based on Bliss Independence, which we find back in our analysis. Irrespective of the origin of the classification, we stress that the labels were provided to us by independent researchers and hence were not biased in any way to favor the Explicit Mean Equation model.

Note that we conduct the research only on combinations of two compounds. Meanwhile, it is shown in Russ and Kishony [2018] that Bliss Independence maintains accuracy when increasing the number of compounds that are combined with each other. Loewe Additivity, however, loses its predictive power for an increasing number of compounds.

The comparison of the parametric method with the lack-of-fit method shows a superiority of the lack-of-fit method. To recall, the motivation behind the parametric approach was the statistical advantages of such an approach. It allows to define an interval around *α* = 0 in which a compound combination can be considered additive. For the lack-of-fit method, such statistical evaluation can not be done directly, but could be performed on the basis of bootstrapping.

Chou and Talalay [1977] measure the interaction effect locally for a fixed ratio of doses of both compounds that are supposed to reach the same effect, say one unit of the first compound causes the same effect as two units of the second compound, which results in the dose combination of 1:2. Along this fixed ratio of doses, they compute the left-hand side of Eq. 3 given the two doses *x*_1_ and *x*_2_ that are assumed to reach a fixed effect *y^*^* together with 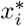 being the dose of compound *i* that reaches the fixed effect alone. For the fixed dose ratio, they run over all expected effects, usually from zero to one. A geometric interpretation of that method is depicted in [Greco et al., 1995, Fig. 7, p. 341]. The resulting values of the left-hand side of Eq. 3 are analyzed graphically: all computed values are plotted versus the expected fixed effect *y*^*^ = [0, 1]. Values higher than one exhibit synergistic behavior, values below one antagonism. This method allows for results that show antagonistic behavior for, say, smaller effects, as well as synergistic behavior for higher effects, or vice versa. That such a behavior of switching from antagonistic behavior in one region to synergistic behavior in another can occur was also shown in Norberg and Wahlström [1988]. With one synergy score, as used throughout this paper, we do not provide such a measure for local antagonism and synergism. Our main motivation in this study is to provide a single synergy score that allows for fast filtering of interesting candidates for more in-depth research. To extend that idea, the standard deviation could be taken into account, as in a *t*-value or Z-score. Additionally, the superior lack-of-fit method is much faster and simpler to implement than the parametric one.

**Figure 7:**
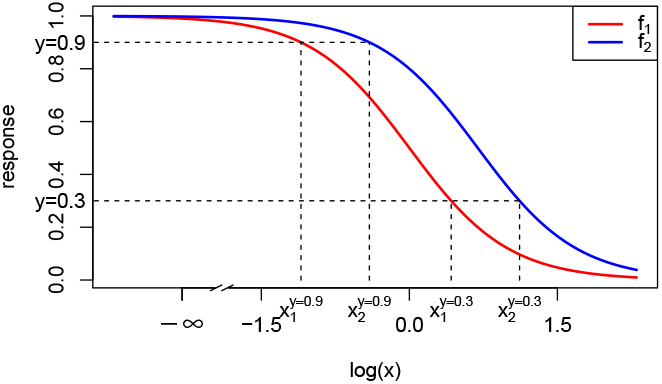
Dose-response curves (red and blue) as Hill curves (Eq. 21). For the exemplary responses of 0.3 and 0.9 the different doses *x*_1_ and *x*_2_ reaching that effect are shown (dashed lines). The dose-response curves differ only in EC_50_ with *e*_1_ = 2 and *e*_2_ = 1. Values of the other parameters are *y*_0_ = 1, *y*_*∞*_ = 0 and *s* = 2. To highlight the sigmoidal shape of a Hill curve in log-space, the logarithmic concentration space is depicted.

Finally, to asses how distinguishable the synergy scores *γ* are, we visualize the synergy scores based on the underlying category (Section 3.2). The synergy scores from the lack-of-fit method can, based on their sign, reliably be categorized as synergistic or antagonistic. For records categorized as non-interactive, the computed synergy scores are positive as well as negative. For the two datasets, we saw different extents of separation between those *γ*-scores, which makes it difficult to generalize the results. All in all, the differentiation from no interaction poses a more difficult task as choosing the threshold is arbitrary.

During the analysis, we observed higher synergy scores when applying the Bliss Independence principle as null reference model. This is due to the more conservative null reference surface as derived from Bliss Independence (see exemplary comparison of isoboles from most of the models discussed here in [Sinzger et al., 2019, Fig. 6]). Due to the synergy scores being relatively high, a differentiation between categories based on the synergy score poses a bigger challenge. There is a strong overlap of synergy scores from all three categories. Additionally, most of the synergy scores *γ*, that are computed with the lack-of-fit method, are above zero. Different ranges of synergy scores for both datasets make it additionally difficult to assess synergy or antagonism for a record based on the unique information of the synergy score.

We want to emphasize the performance benefit of the recently introduced Explicit Mean Equation [Lederer et al., 2018a] over the implicit formulation in form of the General Isobole Equation. On both datasets, it is the overall best performing model when compared to the provided categorizations. The explicit formulation of this additive model was shown to speed up computation by a factor of 250 (see [Lederer et al., 2018a, Fig. S1]). Together with the implementation of the lack-of-fit method, which is easier to implement and a lot faster than the parametric method, this combination of model improvement and method can be of great benefit for the research community.

Although the performance of models and methods are consistent across the two (quite different) datasets considered in this study, reliable comparison of different models and methods would benefit from the availability of drug screening datasets that are available with ground truth labeling.

## Acknowledgments

We thank Bhagwan Yadav for the sharing of the code used for the analysis in Yadav et al. [2015] and Murat Cokol for the sharing of the data and analytical insights from Cokol et al. [2011].

An earlier version of this manuscript has been released as a Pre-Print at [Lederer et al., 2018b].

## A Conditional Dose Response Curves

A common approach for modeling monotonic dose-response curves *f*_*j*_ with *j* ∈ {1, 2} is the Hill curve [Hill, 1910], also referred to as the sigmoid function. The Hill model is, due to its good fit to many sources of data, the most widely applied model for fitting compound responses [Goutelle et al., 2008]. It has a sigmoidal shape with little change for small doses but with a rapid decline in response once a certain threshold is met. For even larger doses the effect asymptotes to a constant maximal effect. Two exemplary Hill curves are depicted in Fig. 7. There are several parameterizations of the Hill curve. We use the following throughout this study to fit conditional responses:

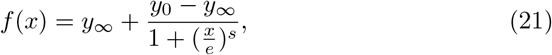

where *y*_0_ is the response at zero dose and *y*_*∞*_ the maximal response of the cells to the compound, *e* the dose concentration reaching half of the maximal response and *s* the steepness of the curve. Eq. 21 is equivalent to the parametrization used in the drc package [Ritz and Strebig, 2016], the so-called four parameter log-logistic model. By our definition of the Hill curve, a positive *s* leads to a descending Hill curve.

## B Data Cleaning, Fitting of Hill Curve and Parameter Estimation for Implicit Models

First, we normalize all records by the measured response at zero dose concentration from both compounds, *y*_0_. Second, we conduct an outlier analysis of the normalized responses by fitting a spline surface and deleting outliers to discard them. Third, we then fit the conditional responses of the cleaned data to Hill curves.

We fit a general additive model (GAM) to the normalized raw data using thin plate splines [Wood, 2017], not transforming the doses in any way. The surfaces of those fitted thin plate splines span the checkerboards of every record and data points with too large absolute residual values are rejected. For fitting the splines we use method gam() of the mgcv-package [Wood, 2011], defining the smooth terms within the gam formulae with the method s(). We set the dimension of the basis, that is used to represent the smooth term to *k* = 30 fixed knots.

The threshold to reject data points is at five times the inter-quantile range of all residuals of a given record. Every data point with an absolute residual above that threshold is discarded. For the Mathews Griner data, this leads to 18 records out of the 466 (less than 4%) where a mean of 1.28 outliers were excluded per record with an overall of 23 data points excluded, which is less than one percent of the overall data. A maximum of 6 outliers was detected once. Similarly, we excluded on average 2.48 data points for the Cokol data on 52 of the total 200 (≤25%) records with a maximum of 13 data points and an overall of 129 data points excluded, which is about 1% of all data points.

To fit the two conditional responses of a record to two Hill functions of the form of Eq. 21 we use the drc package [Ritz et al., 2015]. Unlike other synergy analyses such as [Yadav et al., 2015], the response at zero concentration *y*_0_ is not fixed to 1 but merely constrained to be the same for both response curves. The other Hill parameters, *y_∞_, s* and *e* are fitted for both compounds individually. In case the asymptote parameter *y_∞_* is below zero for any of the two Hill curves, the conditional response of that compound is refitted to a two-parameter model with *y_∞_* set to zero and *y*_0_ kept from the fitting of both compounds together. This is the case for 43 records of the Mathews Griner dataset and 125 records of the Cokol dataset. We exclude records for which any of the Hill curve parameters slope or EC50 are negative (*s* < 0, *e* < 0). This is the case for 187 records for the Mathews Griner dataset (133 records with negative slope *s*, 88 records with negative EC50 value *e*, out of which there are 34 records with negative slope and negative EC50 value), which is roughly 40% of all records. More details follow below.

The *f*_GI_ (*x*_1_, *x*_2_) model is an implicit model for the response *y*. Therefore, a root finder is used to find a response 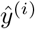 given concentrations 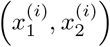 and parameters describing the Hill curves of the conditional responses, Θ = *{y*_0_, *y_∞,j_, e_j_, s_j_*}, We used the standard implementation of a root finder in the R stats package, uniroot() [R Core Team, 2016], which is based on the Brent-Dekker-van Wijngaarden algorithm [Press et al., 2007, Chapter 9]. As convergence criterion we used 1.22 *×* 10^−4^ within a maximum of 1000 iterations.

## B.1 Sensitivity of model performance to inter-quantile range

In a previous version of this article, we cleaned the data to three times the inter-quantile range instead of five. With this smaller inter-quantile range we removed in the Mathews Griner dataset in total 199 data points instead of 23, and in the Cokol dataset 623 instead of 129. The performance of the Mathews Griner dataset for the lack-of-fit method slightly decreased, whereas the overall performance for the Cokol dataset increased (see Appendix E, Fig. 8 and Fig. 9). As a note, approximately the same number of records were excluded for the analysis due to two issues: i) negative slopes of at least one of the conditional dose response curves, or ii) the root-finder for the *f*_CI_ (*x*_1_, *x*_2_|*α*) model not converging (no convergence after 1000 iterations). These issues are independent from data cleaning with three or five times the inter-quantile range (see Appendix D, Table 13 - Table 16).

## B.2 Handling records with negative slope or EC50 values

Roughly 40% (187) of the records of the Mathews Griner dataset were excluded in the study because of a negative slope or EC50 parameter. This is due to a suboptimal choice of doses. We observed two types of sub-optimality: first, the maximal dose can be too small to induce a significant change in response. Due to the noise in measurements, negative slope and EC50 parameters are fitted. This is the case for 34 records. A second type of sub-optimality is observed when the maximal effect is already reached for the second dose (the first dose is always zero). This is the case for the remaining 153 records.

Although we could not fit two reasonable Hill curves to these records, we can still use both methods, lack-of-fit and parametric, to quantify synergy. They both only require two mathematically well-defined conditional response curves. Here, we define a conditional response for cases with negative slope or negative EC50 parameter according to Table 2.

**Table 2:** Response curves for cases where the Hill model fit leads to negative slope or negative EC50 values.

| $s$ | $e$ | $y(x)$ |
| --- | --- | --- |
| $< 0$ | $< 0$ | $y_0 \quad \forall \quad x$ |
| $< 0$ | $> 0$ | $y_\infty$ for $x \neq 0$ |
| $> 0$ | $< 0$ | $y_\infty$ for $x \neq 0$ |

With the above definition for conditional response curves, we investigated the 187 previously excluded records in detail. We computed the lack-of-fit synergy values *γ* for those 187 records and for the entire dataset of 466 records. For two of these records, the *f*_GI_ (*x*_1_, *x*_2_) model did not converge. The Kendall rank correlation coefficient values are given in Appendix D, Table 17. The inclusion of those datasets results in lower Kendall rank correlation coefficients relative to the original analysis. The coefficients decrease by roughly 0.05 when averaged over all models.

## C ROC-analysis

In high-throughput synergy studies, one generally screens for promising candidates that exhibit a synergistic or antagonistic effect. Those promising candidates are then investigated in more detail with genetic assays and other techniques. To determine how well the underlying null reference models result in distinguishable synergy scores, we conduct an ROC analysis (receiver operating characteristic), comparing the estimated synergy scores with the class categorization that is given for both datasets. A standard ROC analysis applies to binary classification, where cases are compared to controls. In this study, we have three classes: synergistic, antagonistic and non-interactive. We therefore compare each class to the combination of the other two, e.g. synergistic as cases versus the antagonistic and non-interactive combined as control. Typically, in ROC analyses, the cases rank higher than the controls. When treating the class antagonistic as case compared to the control synergistic and non-interactive we change all signs of the synergy scores. Therefore, the ranking of synergy scores is reversed and antagonistic synergy scores rank higher. Problems arise when comparing non-interactive cases to the control synergistic and antagonistic as their values should lie between the two control classes. Therefore, the absolute value of the estimated synergy scores is taken, which allows a ranking where the synergy scores of the non-interactive records should rank lower than the other synergy scores. Additionally, we can again multiply all synergy scores with minus one to revert the order of scores such that the cases rank higher.

The AUC values (area under the curve) are reported in Table 5 - Table 8 in Appendix D. For completeness, and based on the critique of Saito and Rehmsmeier [2015] to use PRC-AUC (precision/recall area under the curve) values for imbalanced datasets, the PRC-AUC values are also computed and can be found in Table 9 - Table 12 in Appendix D.

Analogously to the previous section, we depict the AUC values for both datasets in scatter plots (Fig. 10) with AUC values based on the parametric approach depicted on the *x*-axis and those based on the lack-of-fit approach on the *y*-axis. The underlying null reference models are shown by color. The different comparisons, such as synergistic versus non-interactive and antagonistic, are depicted by shape of the plot symbol.

From Fig. 10, the dominance of the lack-of-fit approach over the parametric one is as apparent for the Mathews Griner dataset as from Fig. 2. With regard to the comparison of the different cases, visualized in shape, the AUC values from the comparison of the synergistic cases to the non-interactive and antagonistic controls, score the highest values around 0.9. The comparison of the non-interactive cases to the interactive ones score the lowest.

As the overall highest AUC scores result from the lack-of-fit method, we have a closer look at those for both datasets (Table 6 and Table 8 in Appendix D). For the antagonistic case, the values range around 0.80 for the Mathews Griner dataset and around 0.85 for the Cokol dataset. AUC values of the non-interactive case range around 0.75 for both datasets. The AUC values for the synergistic case for both datasets range around a value of 0.90 with one outlier of 0.75 for the Bliss Independence model on the Mathews Griner dataset. Overall, the lack-of-fit outperforms the parametric method on the Mathews Griner dataset. For the lack-of-fit method, both the *f*_large→small_ (*x*_1_, *x*_2_) and Explicit Mean Equation perform best on the Mathews Griner dataset for synergistic cases, and a clear dominance of *f*_large→small_ (*x*_1_, *x*_2_) over the Explicit Mean Equation for antagonistic and non-interactive cases. On the second dataset, the Cokol dataset, Explicit Mean Equation performs overall best for both methods.

We attribute the differences in performances of methods and models on the two datasets to the differences in the experimental design for these datasets. For the Cokol dataset, all compounds were applied up to their maximal effect dose. In the Mathews Griner dataset, all compounds were applied with the same fixed dose range.

## D Supplementary Tables

**Table 3:** Kendall rank correlation coefficient of Mathews Griner data set.

| model | lack-of-fit | parametric |
| --- | --- | --- |
| $f_{\text{GI}}(x_1, x_2)$ | 0.52 | 0.23 |
| $f_{\text{large} \rightarrow \text{small}}(x_1, x_2)$ | **0.54** | **0.32** |
| $f_{\text{small} \rightarrow \text{large}}(x_1, x_2)$ | 0.48 | 0.19 |
| $f_{\text{mean}}(x_1, x_2)$ | 0.53 | 0.10 |
| $f_{\text{geary}}(x_1, x_2)$ | 0.47 | 0.30 |
| $f_{\text{bliss}}(x_1, x_2)$ | 0.36 | 0.22 |

**Table 4:** Kendall rank correlation coefficient of Cokol data set.

| model | lack-of-fit | parametric |
| --- | --- | --- |
| $f_{\text{GI}}(x_1, x_2)$ | 0.62 | 0.12 |
| $f_{\text{large} \rightarrow \text{small}}(x_1, x_2)$ | 0.61 | 0.50 |
| $f_{\text{small} \rightarrow \text{large}}(x_1, x_2)$ | 0.50 | 0.57 |
| $f_{\text{mean}}(x_1, x_2)$ | **0.67** | **0.64** |
| $f_{\text{geary}}(x_1, x_2)$ | 0.56 | 0.58 |
| $f_{\text{bliss}}(x_1, x_2)$ | 0.56 | 0.16 |

**Table 5:** AUC analysis of parametric method applied to Mathews Griner dataset.

|  | synergistic | non-interactive | antagonistic |
| --- | --- | --- | --- |
| $f_{\text{GI}}(x_1, x_2)$ | 0.67 | 0.63 | 0.62 |
| $f_{\text{large} \rightarrow \text{small}}(x_1, x_2)$ | **0.73** | **0.68** | **0.72** |
| $f_{\text{small} \rightarrow \text{large}}(x_1, x_2)$ | 0.68 | 0.53 | 0.53 |
| $f_{\text{mean}}(x_1, x_2)$ | 0.62 | **0.68** | 0.46 |
| $f_{\text{geary}}(x_1, x_2)$ | 0.70 | 0.57 | 0.60 |
| $f_{\text{bliss}}(x_1, x_2)$ | 0.66 | 0.45 | 0.62 |

**Table 6:** AUC analysis on lack-of-fit method applied to Mathews Griner dataset.

|  | synergistic | non-interactive | antagonistic |
| --- | --- | --- | --- |
| $f_{\text{GI}}(x_1, x_2)$ | 0.88 | 0.77 | 0.81 |
| $f_{\text{large} \rightarrow \text{small}}(x_1, x_2)$ | **0.89** | **0.78** | **0.84** |
| $f_{\text{small} \rightarrow \text{large}}(x_1, x_2)$ | 0.86 | 0.68 | 0.78 |
| $f_{\text{mean}}(x_1, x_2)$ | **0.89** | 0.75 | 0.82 |
| $f_{\text{geary}}(x_1, x_2)$ | 0.85 | 0.69 | 0.78 |
| $f_{\text{bliss}}(x_1, x_2)$ | 0.75 | 0.60 | 0.76 |

**Table 7:** AUC analysis on parametric method applied to Cokol dataset.

|  | synergistic | non-interactive | antagonistic |
| --- | --- | --- | --- |
| $f_{\text{GI}}(x_1, x_2)$ | 0.62 | 0.62 | 0.55 |
| $f_{\text{large} \rightarrow \text{small}}(x_1, x_2)$ | 0.80 | 0.64 | 0.84 |
| $f_{\text{small} \rightarrow \text{large}}(x_1, x_2)$ | 0.86 | 0.64 | 0.89 |
| $f_{\text{mean}}(x_1, x_2)$ | **0.89** | **0.74** | **0.93** |
| $f_{\text{geary}}(x_1, x_2)$ | 0.80 | **0.74** | 0.89 |
| $f_{\text{bliss}}(x_1, x_2)$ | 0.62 | 0.55 | 0.59 |

**Table 8:**
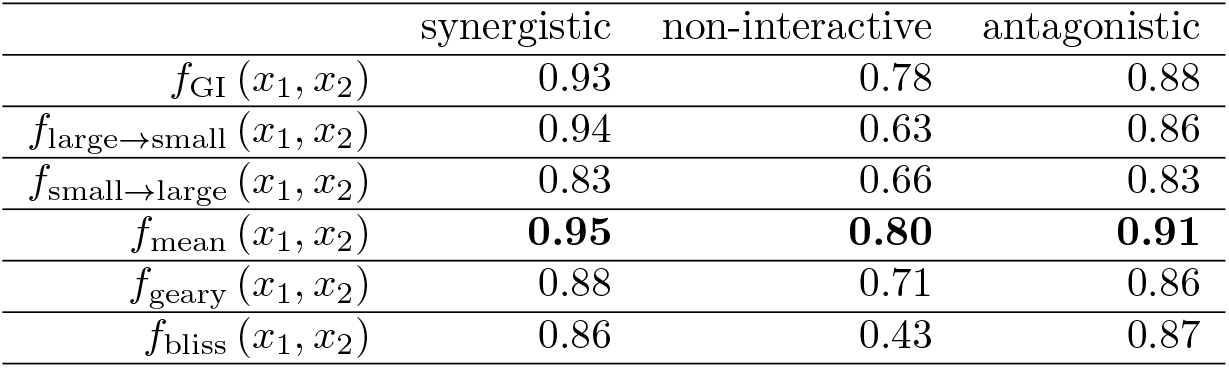
AUC analysis on lack-of-fit method applied to Cokol dataset.

|  | synergistic | non-interactive | antagonistic |
| --- | --- | --- | --- |
| $f_{\text{GI}}(x_1, x_2)$ | 0.93 | 0.78 | 0.88 |
| $f_{\text{large} \rightarrow \text{small}}(x_1, x_2)$ | 0.94 | 0.63 | 0.86 |
| $f_{\text{small} \rightarrow \text{large}}(x_1, x_2)$ | 0.83 | 0.66 | 0.83 |
| $f_{\text{mean}}(x_1, x_2)$ | **0.95** | **0.80** | **0.91** |
| $f_{\text{geary}}(x_1, x_2)$ | 0.88 | 0.71 | 0.86 |
| $f_{\text{bliss}}(x_1, x_2)$ | 0.86 | 0.43 | 0.87 |

**Table 9:**
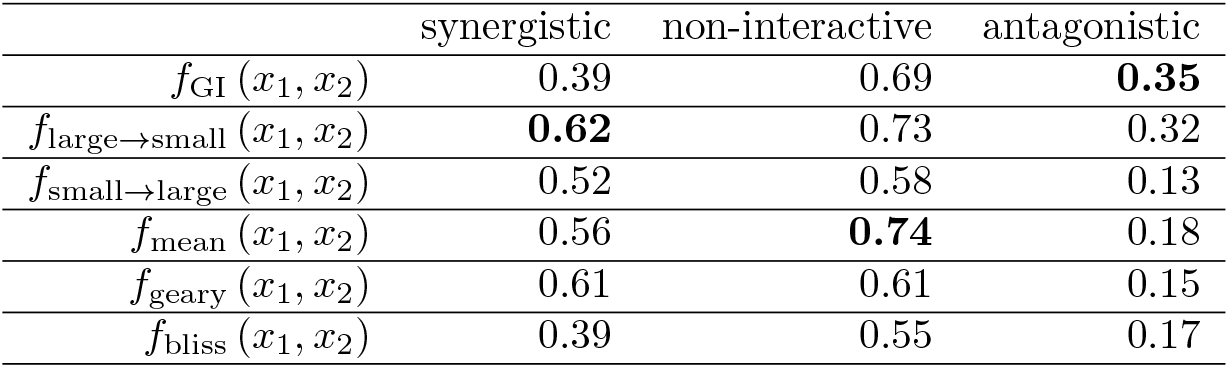
PRC-AUC analysis on parametric method applied to Mathews Griner dataset.

|  | synergistic | non-interactive | antagonistic |
| --- | --- | --- | --- |
| $f_{\text{GI}}(x_1, x_2)$ | 0.39 | 0.69 | **0.35** |
| $f_{\text{large} \rightarrow \text{small}}(x_1, x_2)$ | **0.62** | 0.73 | 0.32 |
| $f_{\text{small} \rightarrow \text{large}}(x_1, x_2)$ | 0.52 | 0.58 | 0.13 |
| $f_{\text{mean}}(x_1, x_2)$ | 0.56 | **0.74** | 0.18 |
| $f_{\text{geary}}(x_1, x_2)$ | 0.61 | 0.61 | 0.15 |
| $f_{\text{bliss}}(x_1, x_2)$ | 0.39 | 0.55 | 0.17 |

**Table 10:** PRC-AUC analysis on lack-of-fit method applied to Mathews Griner dataset.

|  | synergistic | non-interactive | antagonistic |
| --- | --- | --- | --- |
| $f_{GI}(x_1, x_2)$ | 0.78 | 0.75 | 0.48 |
| $f_{large \rightarrow small}(x_1, x_2)$ | **0.80** | 0.74 | **0.55** |
| $f_{small \rightarrow large}(x_1, x_2)$ | 0.72 | 0.69 | 0.33 |
| $f_{mean}(x_1, x_2)$ | 0.78 | **0.76** | 0.42 |
| $f_{geary}(x_1, x_2)$ | 0.71 | 0.70 | 0.35 |
| $f_{bliss}(x_1, x_2)$ | 0.52 | 0.64 | 0.39 |

**Table 11:** PRC-AUC analysis on parametric method applied to Cokol dataset.

|  | synergistic | non-interactive | antagonistic |
| --- | --- | --- | --- |
| $f_{GI}(x_1, x_2)$ | 0.30 | 0.57 | 0.51 |
| $f_{large \rightarrow small}(x_1, x_2)$ | 0.56 | 0.56 | 0.65 |
| $f_{small \rightarrow large}(x_1, x_2)$ | 0.62 | 0.52 | 0.83 |
| $f_{mean}(x_1, x_2)$ | **0.71** | 0.55 | **0.89** |
| $f_{geary}(x_1, x_2)$ | 0.60 | **0.58** | 0.82 |
| $f_{bliss}(x_1, x_2)$ | 0.30 | 0.45 | 0.50 |

**Table 12:** PRC-AUC analysis on lack-of-fit method applied to Cokol dataset.

|  | synergistic | non-interactive | antagonistic |
| --- | --- | --- | --- |
| $f_{GI}(x_1, x_2)$ | 0.84 | 0.65 | 0.83 |
| $f_{large \rightarrow small}(x_1, x_2)$ | **0.87** | 0.46 | 0.72 |
| $f_{small \rightarrow large}(x_1, x_2)$ | 0.63 | 0.56 | 0.72 |
| $f_{mean}(x_1, x_2)$ | **0.87** | **0.66** | **0.86** |
| $f_{geary}(x_1, x_2)$ | 0.75 | 0.60 | 0.77 |
| $f_{bliss}(x_1, x_2)$ | 0.76 | 0.35 | 0.70 |

**Table 13:** \# of excluded records from parametric and lack-of-fit method applied to the Mathews Griner dataset with a threshold of three times the inter-quantile range.

|  | synergistic | antagonistic | non-interactive | total |
| --- | --- | --- | --- | --- |
| parametric | 19 | 15 | 48 | 82 |
| lack-of-fit | 34 | 59 | 93 | 186 |
| both | 19 | 16 | 49 | 84 |

**Table 14:**
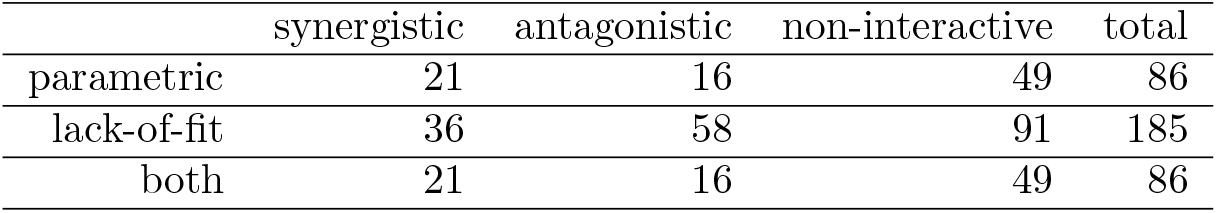
\# of excluded records from the parametric and lack-of-fit method applied to the Mathews Griner dataset with cleaned data of a threshold of five times the inter-quantile range.

|  | synergistic | antagonistic | non-interactive | total |
| --- | --- | --- | --- | --- |
| parametric | 21 | 16 | 49 | 86 |
| lack-of-fit | 36 | 58 | 91 | 185 |
| both | 21 | 16 | 49 | 86 |

**Table 15:**
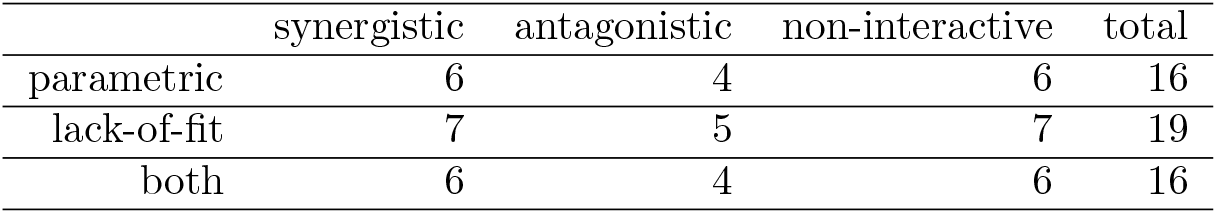
\# of excluded records from parametric and lack-of-fit method applied to the Cokol dataset with cleaned data of a threshold of three times the inter-quantile range.

**Table 16:** \# of excluded records from parametric and lack-of-fit method applied to Cokol dataset with cleaned data of a threshold of five times the inter-quantile range.

|  | synergistic | antagonistic | non-interactive | total |
| --- | --- | --- | --- | --- |
| parametric | 3 | 2 | 4 | 9 |
| lack-of-fit | 3 | 3 | 4 | 10 |
| both | 3 | 2 | 4 | 9 |

**Table 17:** Kendall rank correlation coefficients with recomputed conditional response curves according to Table 2 on 185 records with negative slope or EC50 value (left) and on entire dataset with 464 records (right).

| model | subset of<br>185 records | entire set of<br>464 records | original analysis on<br>279 records |
| --- | --- | --- | --- |
| $f_{GI}(x_1, x_2)$ | 0.37 | **0.47** | 0.52 |
| $f_{large \rightarrow small}(x_1, x_2)$ | 0.34 | 0.43 | **0.54** |
| $f_{small \rightarrow large}(x_1, x_2)$ | **0.42** | 0.46 | 0.48 |
| $f_{mean}(x_1, x_2)$ | 0.37 | 0.43 | 0.53 |
| $f_{geary}(x_1, x_2)$ | 0.35 | 0.39 | 0.47 |
| $f_{bliss}(x_1, x_2)$ | 0.27 | 0.34 | 0.36 |

## E Supplementary Figures

**Figure 8:**
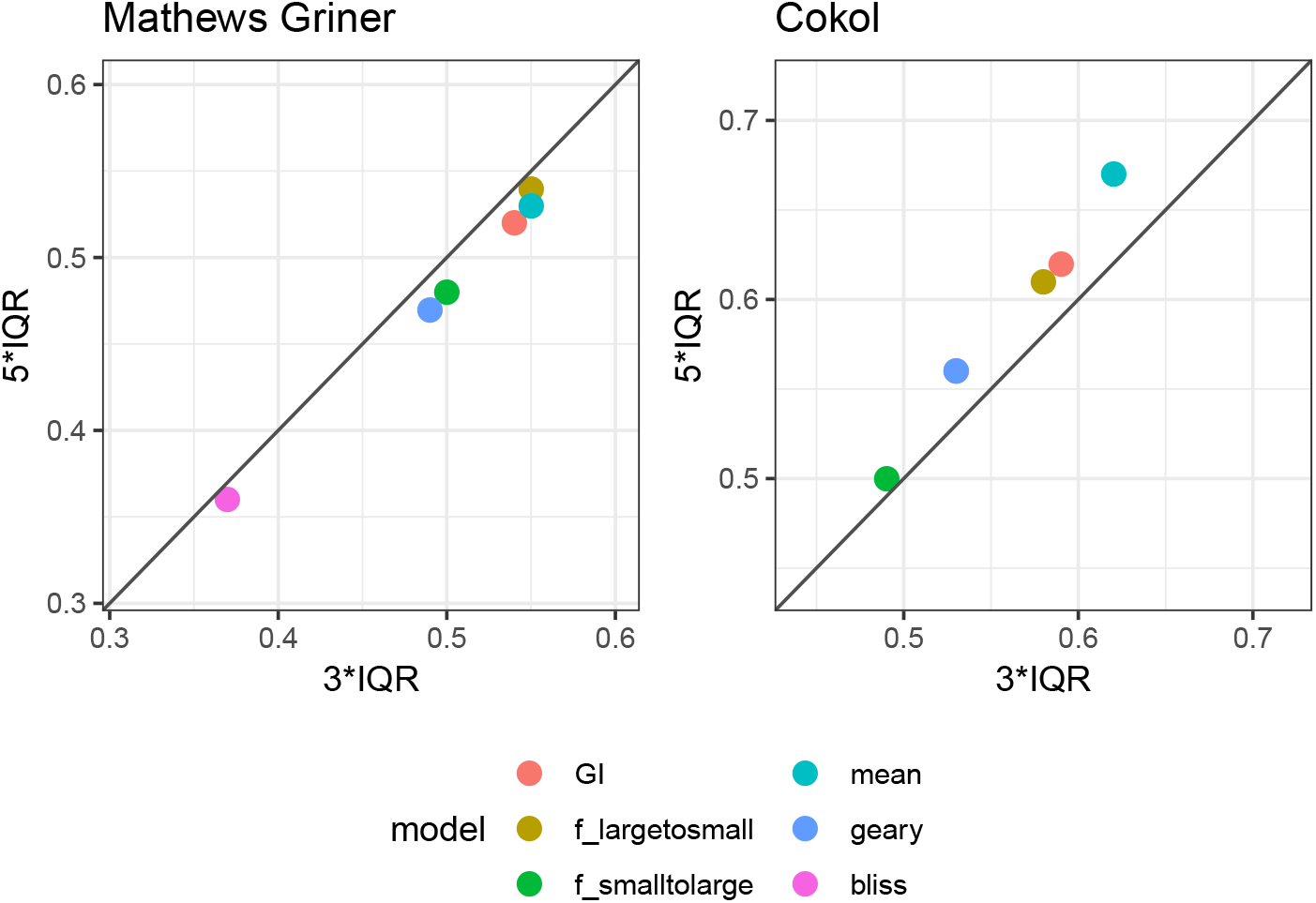
Scatter plot of Kendall rank correlation coefficient for both datasets, Mathews Griner (left) and Cokol (right), comparing the performance of the lack-of-fit method for different data-cleaning thresholds. The Kendall rank correlation coefficient values resulting from the cleaned data with three times the inter-quantile range are plotted on the *x*-axis and those from the data cleaned with a threshold of five times inter-quantile range are plotted on the *y*-axis. Each model is depicted in a different colour. To guide the eye, the diagonal is plotted.

**Figure 9:**
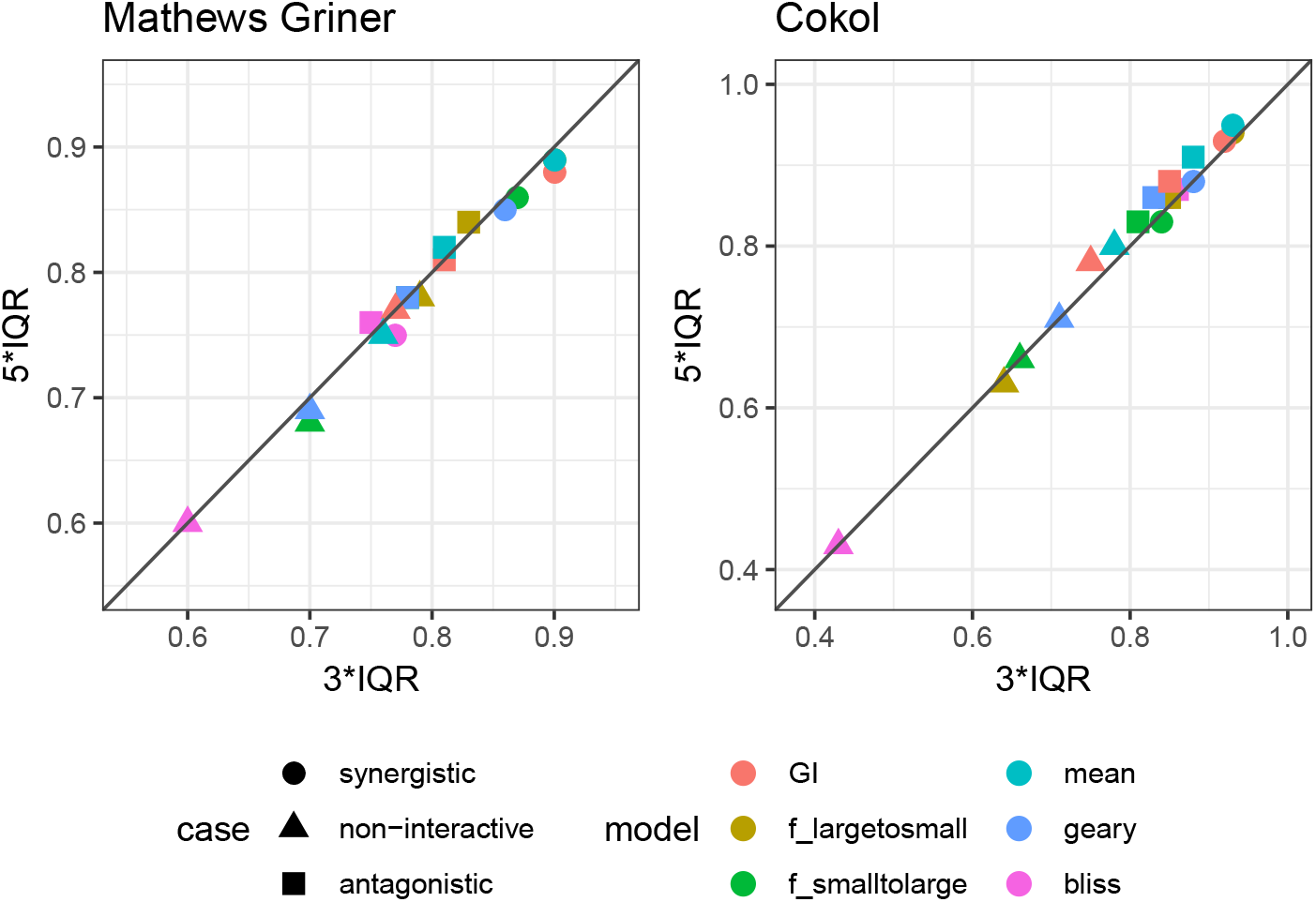
Scatter plot of ROC analysis for both datasets, Mathews Griner (left) and Cokol (right), comparing the performance of the lack-of-fit method for different data-cleaning thresholds. The AUC values resulting from the cleaned data with three times the inter-quantile range are plotted on the *x*-axis and those from the data cleaned with a five times inter-quantile range are plotted on the *y*-axis. AUC values from different models are shown in different colors. AUC values comparing the different categories are depicted in different shapes, where the naming of the shape represents the category that is compared to the remaining two. To guide the eye, the diagonal is plotted. The more a datapoint is above the diagonal, the better the performance of the data cleaned with a threshold of five times the inter-quantile range, and vice versa.

**Figure 10:**
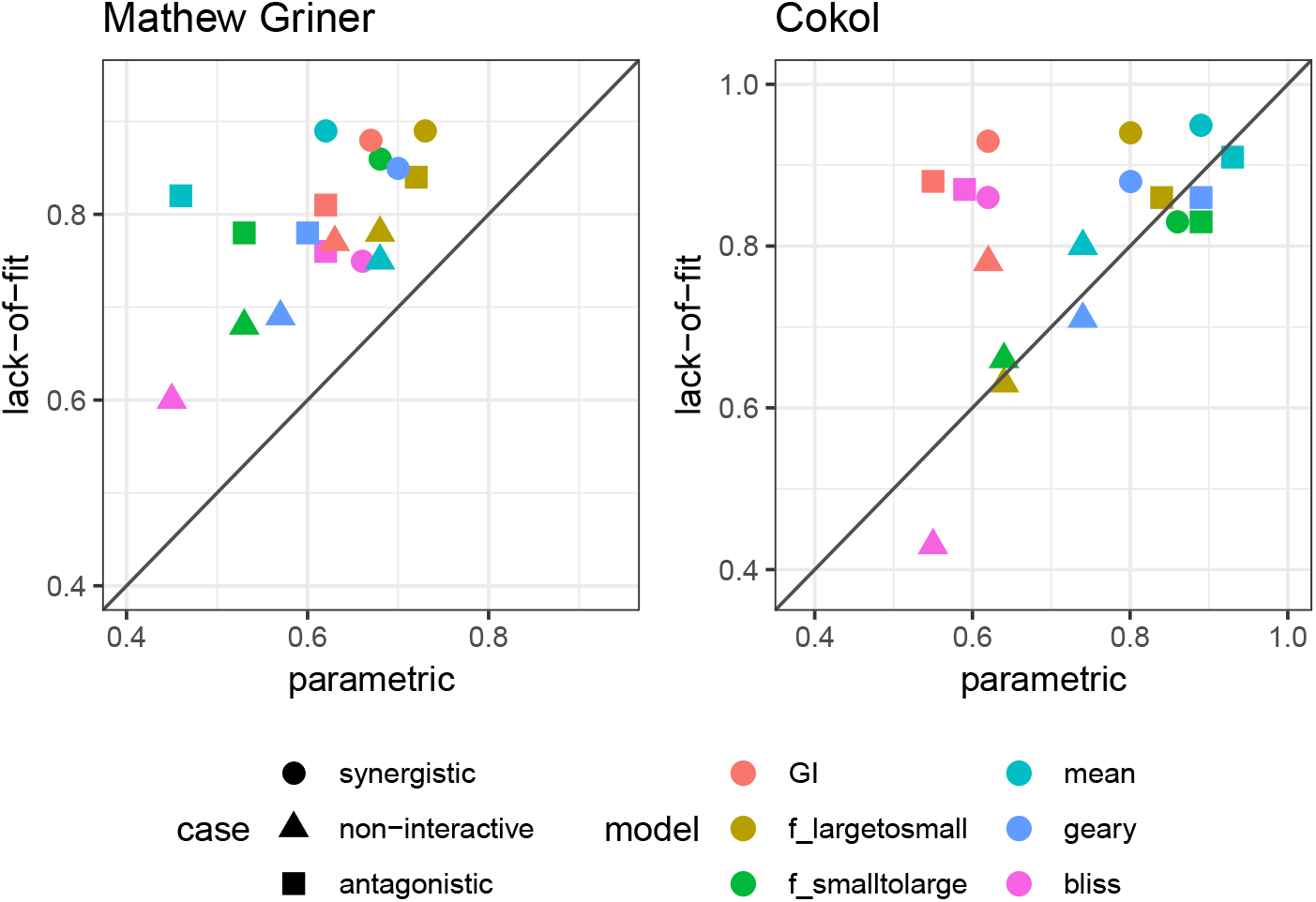
Scatter plot of ROC values for both datasets, Mathews Griner (left) and Cokol (right). The ROC values resulting from the parametric approach are plotted on the *x*-axis and those from the lack-of-fit approache are plotted on the *y*-axis. Each model is depicted in a different color. The three different comparisons, of one case versus the remaining two, are depicted in different shapes. To guide the eye, the diagonal is plotted. If a data point is above the diagonal, the ROC value from the lack-of-fit method is higher than that from the parametric method, and vice versa. Except for the non-interactive comparison of the Bliss Independence model, the synergy scores *γ* from the lack-of-fit method always result in higher ROC values than those computed based on the synergy scores *α* from the parametric method.

**Figure 11:**
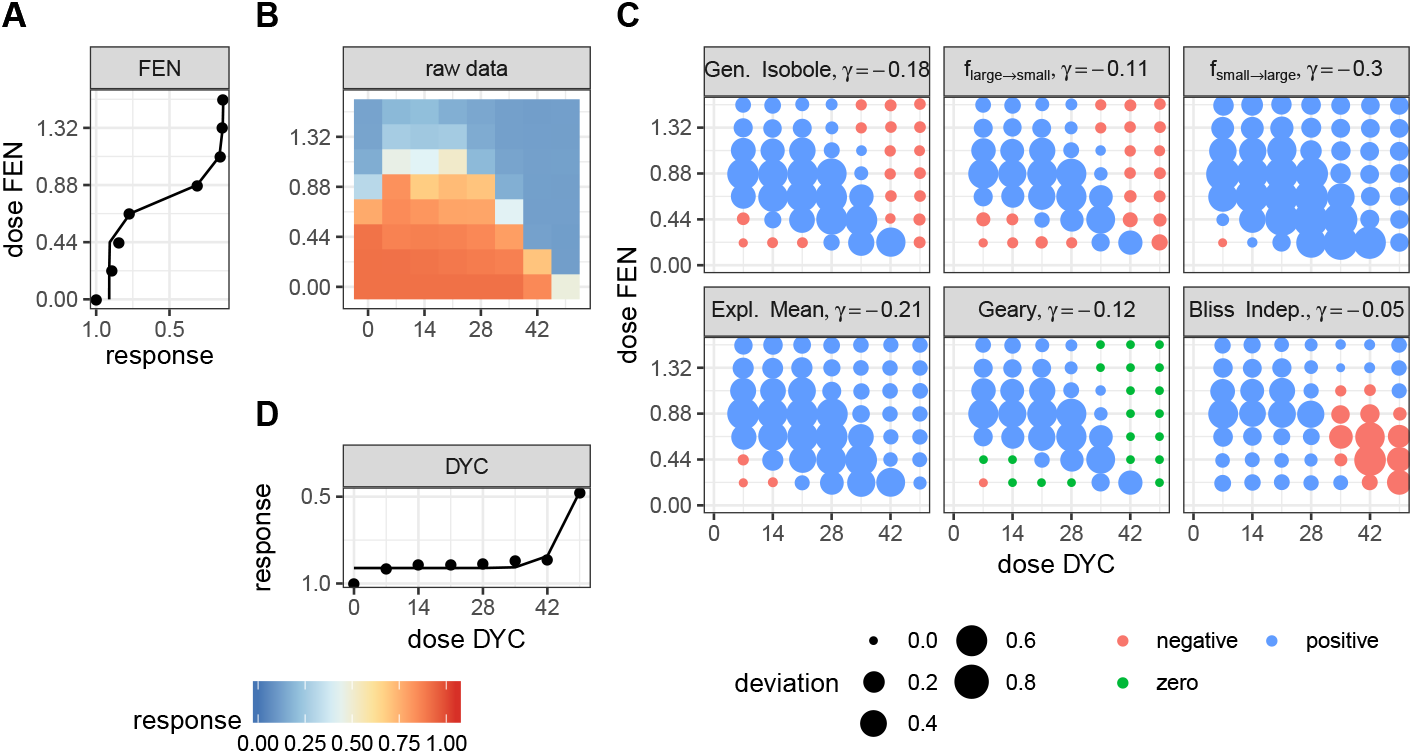
Description of the analysis steps of the lack-of-fit method for the compound pair FEN and DYC from the Cokol dataset. This compound pair is categorized as antagonistic according to [Cokol et al., 2011]. The raw response data of the record is depicted in (B). The response data normalized by the read at zero dose concentration (lower left). In (B) the degree of relative cell growth is colored from high to low values in red to blue. Step 1: compute Hill curves for conditional responses: Based on the raw reads of the single dose responses (lower and left outer edges) fit a Hill curve to the conditional responses. The fitted Hill curves shown in (A) and (D) with original raw data shown as points. Step 2: compute expected non-interactive response for all six models: not shown. Step 3: compute difference between measured data (C) and expected data from all six null reference models: shown in (C). The direction of difference is shown by color (red for negative and blue for positive, green for zero). The larger the degree of difference, the larger the bullet, and vice versa. Step 4: compute integral *γ* over the differences: Over all those bullets, we then compute the integral, which gives the synergy score *γ*. For every model, the synergy score *γ* is depicted in the title of each matrix in (C).

**Figure 12:**
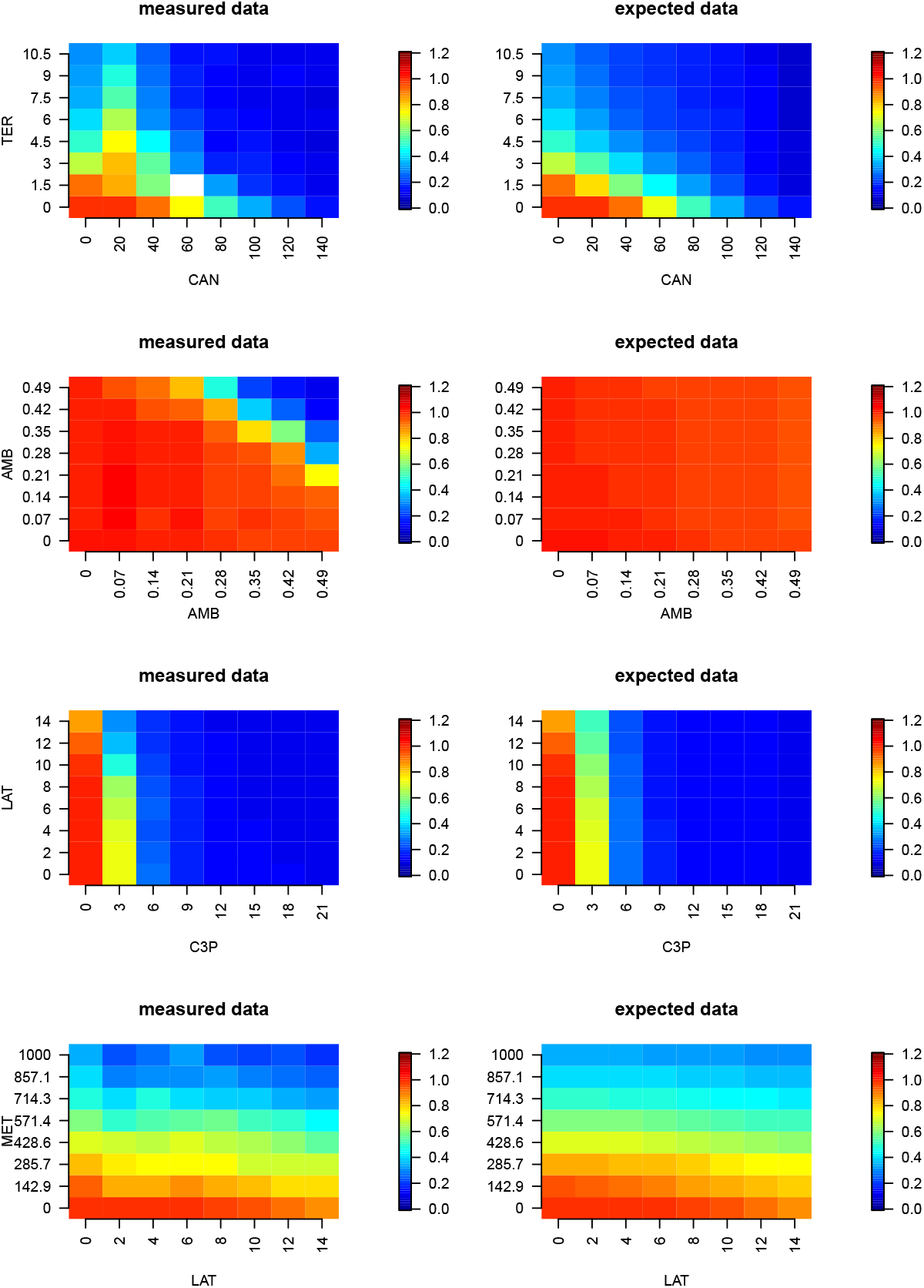
Raw responses (left) and expected responses from the Explicit Mean Equation model (right) of the four records from the Cokol dataset, for which the General Isobole Equation and Explicit Mean Equation gave synergy scores of opposite sign to the orignal categorization. More details on some parameters of the Hill curves can be found in Fig. 6.

